# High-resolution *In-situ* Structures of Mammalian Mitochondrial Respiratory Supercomplexes in Reaction within Native Mitochondria

**DOI:** 10.1101/2024.04.02.587796

**Authors:** Wan Zheng, Pengxin Chai, Jiapeng Zhu, Kai Zhang

## Abstract

Mitochondria play a pivotal role in ATP energy production through oxidative phosphorylation, which occurs within the inner membrane via a series of respiratory complexes. Despite extensive *in-vitro* structural studies, revealing the atomic details of their molecular mechanisms in physiological states remains a major challenge, primarily because of the loss of the native environment during purification. Here, we directly image porcine mitochondria using an *in-situ* cryo-electron microscopy approach. This enables us to determine the structures of various high-order assemblies of respiratory supercomplexes in their native states, achieving up to 1.8-Å local resolution. We identify four major supercomplex organizations: I1III2IV1, I1III2IV2, I2III2IV2, and I2III4IV2, which can potentially expand into higher-order arrays on the inner membranes. The formation of these diverse supercomplexes is largely contributed by ‘protein-lipids-protein’ interactions, which in turn dramatically impact the local geometry of the surrounding membranes. Our *in-situ* structures also capture numerous reactive intermediates within these respiratory supercomplexes, shedding light on the dynamic processes of the ubiquinone/ubiquinol exchange mechanism in complex I and the Q-cycle in complex III. By comparing supercomplex structures from mitochondria treated under distinct conditions, we elucidate how conformational changes and ligand binding states interplay between complexes I and III in response to environmental redox alterations. Our approach, by preserving the native membrane environment, enables structural studies of mitochondrial respiratory supercomplexes in reaction at high resolution across multiple scales, spanning from atomic-level details to the broader subcellular context.

## Introduction

Mitochondria stand central to energy production in eukaryotic cells^1–3^. Mitochondrial dysfunctions correlate with a broad array of severe ailments, including metabolic, cardiovascular, neurodegenerative, and neuromuscular diseases^4–6^. Eukaryotic mitochondria comprise over a thousand proteins, which dynamically assemble into various forms of complexes^7,8^. A key group among these includes the respiratory chain supercomplexes, primarily consisting of complexes I, III, and IV (CI, CIII, and CIV) in varying stoichiometries, serving as the minimal functional units for NADH:O2 oxidoreduction^9^. Recent studies in cryo-electron microscopy (cryo-EM) have illuminated structures of several mammalian respiratory complexes^10–14^ and supercomplexes^15–18^, derived from *in-vitro* purified protein, revealing their subunit compositions, conformational dynamics, and ligand-binding sites. Mechanistically, these structures have shed light on the electron transfer pathways, proton-pumping mechanisms, and regulatory elements^10–18^. Cryo-EM has also provided insights into possible assemblies within mitochondria and hypothetical functional roles of these respiratory supercomplexes^15–19^. However, traditional *in-vitro* approaches result in a loss of the native environment, which poses a significant challenge in elucidating the actual assemblies and molecular mechanisms under physiological conditions.

Here, we report *in-situ* structures of porcine respiratory supercomplexes, determined directly from imaging porcine mitochondria by single-particle analysis combined with cryo-electron tomography (Fig. 1, Extended Data Fig. 1-7, Supplementary Video 1 and Supplementary Information Table 1-7). Our structures, with an average resolution of ∼2.5 Å and local resolution up to 1.8 Å in the best regions, enable in-depth study of mitochondrial respiratory supercomplexes in their native states. With this resolution, we resolve numerous reactive intermediates within these supercomplexes, and determine structures of the four major types of supercomplex organizations.

**Fig. 1.**
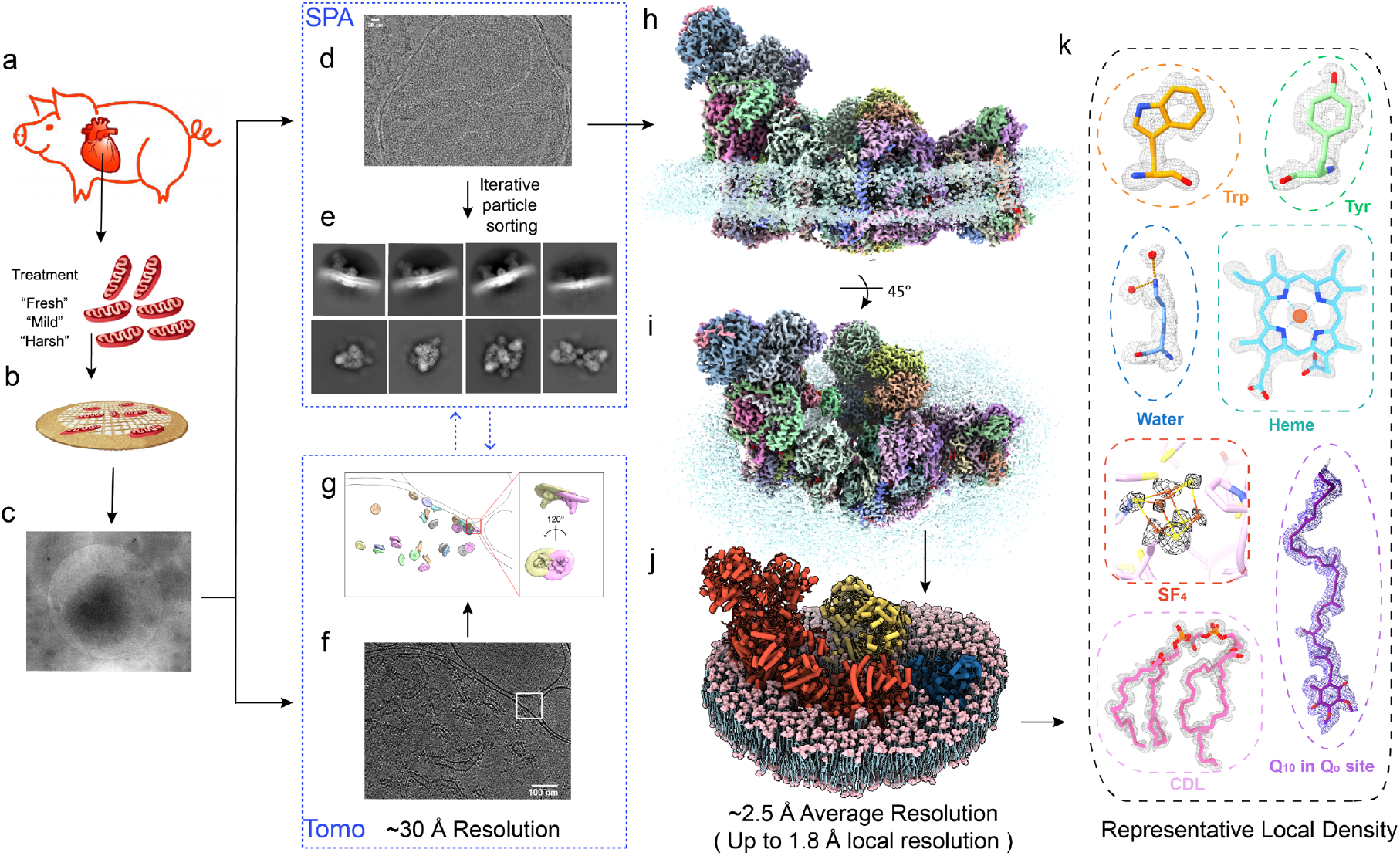
| *In-situ* Single-Particle Cryo-EM and Cryo-ET Analysis of Mammalian Mitochondrial Respiratory Supercomplexes. **a-c,** Mitochondria were extracted from porcine hearts treated under different conditions (fresh, mild and harsh) and were directly frozen onto cryo-EM grids for subsequent structural studies. **d,** Representative single-particle cryo-EM micrograph of mitochondria. **e,** Representative 2D class averages showing different types of supercomplexes after 3D classification. **f, g,** Representative subvolume averages and corresponding slices of reconstructed tomograms. **h, i,** Side and top views of a representative high-resolution map of the supercomplex in the native mitochondrial inner membrane. **j,** A molecular model of the high-resolution supercomplex with the surrounding membrane built. **k,** High-resolution features shown by representative density of amino acid residues and endogenous ligands.

Our structures show distinct classes with different density patterns for endogenous ubiquinone/ubiquinol (Q/QH2) and its interacting residues, providing new structural insights into Q10 exchange dynamics within CI. We also capture multiple states of Q10 bound to CIII’s Qo sites, along with positional shifts in the Rieske domain. Our high-resolution maps unambiguously show complex hydrogen-bond networks, offering detailed perspectives of the proton transfer and the Q-cycle mechanism.

To assess the impact of pathogenic conditions on these complexes, we subject porcine hearts to various treatments mimicking different levels of myocardial ischemia, which affect the distribution of reactive states in the respiratory complexes without compromising the resolution of the cryo-EM maps. Overall, our *in-situ* approach enables the potential for investigating the impacts of diverse mitochondrial diseases and pharmacological treatments by determining reactive protein structures under physiological conditions within mitochondria.

### Structures of supercomplexes with different compositions

Three-dimensional (3D) reconstruction unveils diverse forms of supercomplexes with distinct compositions, including four dominant types: I1III2IV1 (type-A), I1III2IV2 (type-B), I2III2IV2 (type-O), and I2III4IV2 (type-X) (Fig. 2a-d and Extended Data Fig. 3-7). Type-A and -O are similar to previously reported *in-vitro* structures^10–14^, while type-B and -X represents two novel forms not observed *in vitro*. Our cross-classification results indicate that the type-A supercomplex (Fig. 1h-j) is the most abundant form, which is determined at 1.8-2.4 Å resolutions in most CI and CIII regions (Extended Data Fig. 3 and 8), and ∼2.9 Å in CIV (Extended Data Fig. 3). The high-quality maps, as demonstrated by discernible ’holes’ in the side chains of aromatic residues, the 5-element pyrrole rings in heme, and to pinpoint the precise positions of S/Fe atoms in SF4 and the Fe atom in heme, enable us to build accurate atomic models of this form (Fig. 1k, Extended Data Fig. 8 and Supplementary Video 2). In type-A supercomplex, we model 5,297 water molecules into the structure, including those that likely play central roles in proton transfer across the membrane. Furthermore, we build 204 structured and associated lipids (Fig. 2i-k, Extended Data Fig. 9 and Supplementary Video 3), which contribute to stabilizing the protein structure, enhancing the stability of the supercomplexes by facilitating essential protein-lipid-protein interactions (Fig. 2i-k), creating a hydrophobic environment in the Q-binding sites (Supplementary Video 3) and participating in the hydrogen-bond network with their polar heads. Surrounding the proteins, we directly visualize mitochondrial inner membranes composed of more dynamic lipids and build the atomic models (Fig. 2i-k, Extended Data Fig. 9 and Supplementary Video 3). While the overall architecture resembles a previously reported structure using *in-vitro* purified protein^15^, our high-resolution *in-situ* structure shows substantial differences at the interaction interfaces among complexes I, III2, and IV (Extended Data Fig. 10e, f). The membrane enveloping the supercomplex exhibits noticeable curvature, varying across different regions. Specifically, the membrane at the CI heel bends toward the mitochondrial matrix, whereas the CIII2 region displays an opposite curvature pattern (Fig. 2e-h, Extended Data Fig. 10a-d, and Supplementary Video 4).

**Fig. 2.**
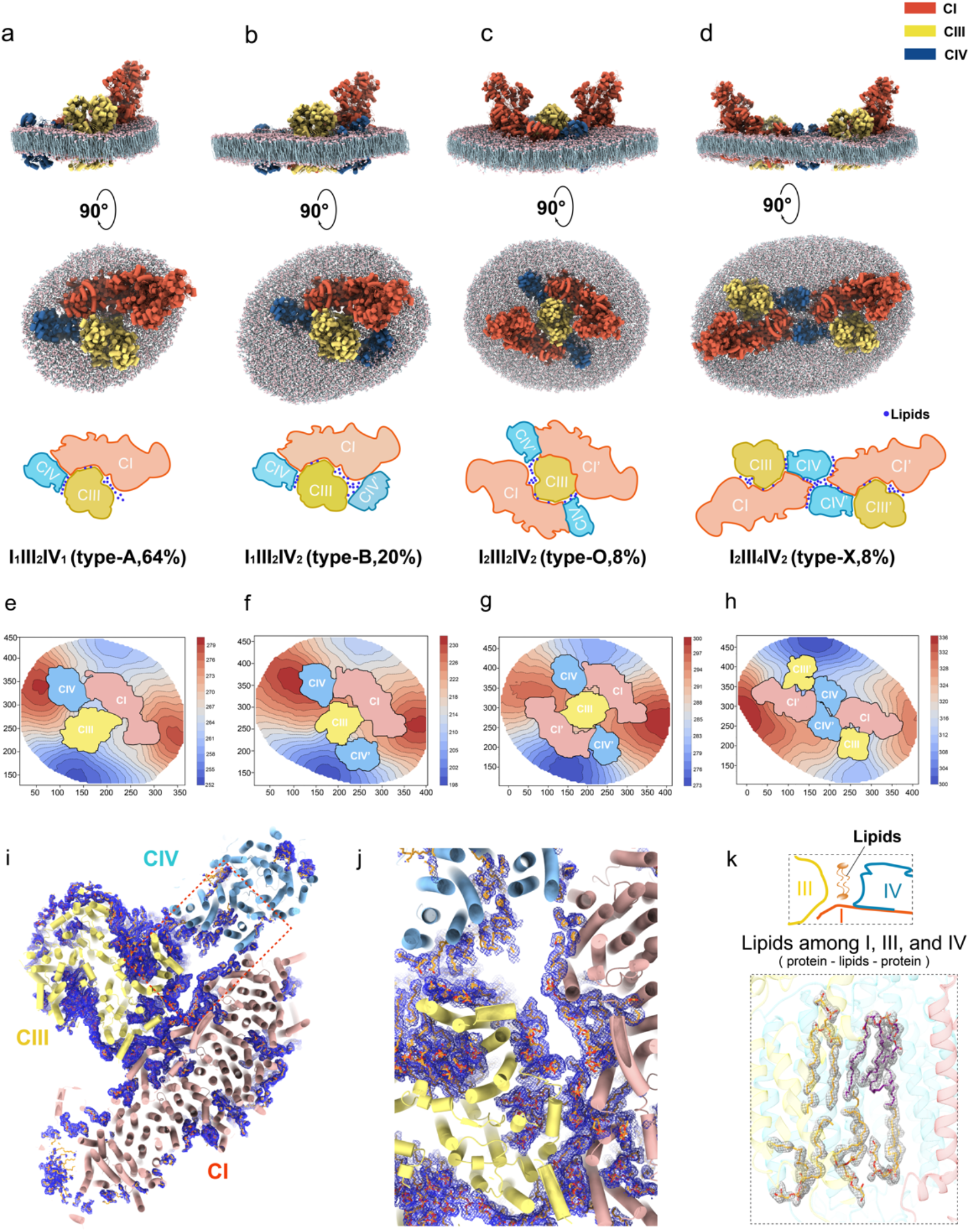
| Architecture, membrane curvature and interaction interfaces of the four types of respiratory supercomplexes. **a-d**, Top and side views of the supercomplexes I₁III₂IV₁, I₁III₂IV₂, I₂III₂IV₂, and I₂III₄IV₂ with models of the surrounding membranes built. These views highlight the impact of different supercomplex compositions on local membrane curvature. **e-h,** Contour maps of the native membrane around the four major types of supercomplexes, viewed from the mitochondrial matrix side. Red and blue indicate high and low altitudes, respectively. These gradients clearly demonstrate the common feature that the membranes surrounding the CI heel and CIII regions are convex and concave, respectively. They also show distinct local curvature differences among the four types. **i, j** Interaction interfaces among CI (light red), CIII (light yellow) and CIV (light blue) in type-A supercomplex. Blue mesh represents the density maps of lipid molecules filling the interstitial space among these complexes. **k,** Representative local density maps and the atomic modes of lipids built in the interface between CIII and CIV.

Our high-resolution density maps of native respiratory supercomplexes allow for accurate analysis of the interaction interfaces. In type-A supercomplex (Fig. 2a), CI-CIII2 interactions are mediated through specific contacts between the subunits NDUFA11^I^ and UQCRQ^III^, as well as NDUFB9^I^, NDUFB4^I^, and UQCRC1^III^. For CI-CIV interactions, the key participating subunits are NDUFB3^I^, ND5^I^, and COX7A^IV^ (Extended Data Fig. 10e, f). One striking feature of the *in-situ* structure is that the interstitial space among CI, CIII and CIV is populated by lipids that mediate the complex interactions. In particular, direct protein-protein interactions between CIII2 and CIV are not observed in our structures (Fig. 2k).

In contrast to type-A, the type-B supercomplex (Fig. 2b) incorporates a second CIV(CIV’) situated between CIII2 and the hydrophilic arms of CI. Unlike CIII2, which exhibits 2-fold symmetry, this additional CIV in the type-B supercomplex is not 2-fold symmetric to the first CIV relative to the symmetry axis of CIII. Instead, it displays a ∼60-degree rotation (Extended Data Fig. 11a, b), partially enclosing the lipid bilayer between the Q-binding pockets in CI and CIII, forming an architecture that could confine the free diffusion of Q10 and facilitate the Q-channeling. In type-B supercomplex, CIV’ establishes novel interactions with CI via the NDUFA1^I^ and COX5A^IV’^ subunits, and with CIII2 via the UQCR10^III^ and COX6A^IV’^ subunits (Extended Data Fig. 10f).

Additionally, two higher-order supercomplexes are determined via a multi-level cross-classification approach we developed. The I2III2IV2 (Fig. 2c) complex at 2.6-3.3 Å resolutions (Extended Data Fig. 5), designated as type-O due to its overall shape, resembles the human mitochondrial megacomplex^15^, and can be considered a pseudo-C2 symmetric expansion of type-A, sharing the CIII dimer (Extended Data Fig. 11c). In contrast to the detergent-purified type-O supercomplex, we observed that the native type-O supercomplex exhibits pseudo-C2 rather than strict C2 symmetry, with the surrounding membrane bending toward the mitochondrial matrix. This observation is not surprising when considering the highly curved membrane structures of the cristae.

The novel supercomplex I2III4IV2 (Fig. 2d), termed type-X owing to its chromosome-like appearance, is formed by two type-A supercomplexes aligned in a head-to-head manner along a pseudo-2-fold axis (Extended Data Fig. 11d). Intriguingly, no canonical strong protein-protein interactions are evident at the dimerization interfaces between the two type-A supercomplexes, except for potential weak contacts between helices NDUFB3^I^ and COX4^IV^. The dimerization is primarily mediated by ’protein-lipids-protein’ interactions in our structure, as evidenced by the numerous lipid molecules filling the interfacial spaces (Supplementary Video 3).

Notably, the formation of the supercomplexes dramatically influences the local curvature of the surrounding membranes (Fig. 2e-h, Extended Data Fig. 10a-d and Supplementary Video 4). Vice versa, we speculate the membrane geometry in turn affects the overall arrangement, distribution and conformation of these supercomplexes, which is worthy of further investigation. In addition, we detect potential extensions of respiratory supercomplexes into high-order arrays in the mitochondrial inner membranes.

### Multiple ubiquinone/ubiquinol-binding states of complex I captured in reaction

CI orchestrates electron transfer from NADH to ubiquinone and concurrently translocases protons across the inner mitochondrial membrane^20^. The Q-binding site in CI features an unusually long, heterogeneous channel structure^21–23^. The entry, exit, and interactions of Q10 within this channel are central yet unresolved questions in understanding CI’s molecular mechanisms. *In-vitro* methods have been limited in detailing this dynamic process at the atomic level.

We find five major Q-binding states (Fig. 3), which include the apo state and a previously described Q-bound state^21,24^ (State-1), along with three novel Q-bound states (State 2-4) (Fig. 3b-e). The apo-state displays only noisy densities in the Q-channel. State-1 resembles the active form of bovine CI (PDB code: 7QSK^21^) (Fig. 3f), with a Q10 molecule fully occupying the Q-binding site (Fig. 3c). In this state, the 1-carbonyl group of Q10 is situated ∼20 Å (Fig. 3b) above the membrane surface (M-distance) and 41 Å (Fig. 3d) from the channel entrance (E-distance). The three other states show not only conformational differences in the Q10 headgroup and the key residue H59^NDUFS^^2^, but also long-range structural changes away from the Q10 headgroup and along the whole Q10. In State-2, the Q headgroup is angled away from the FeS cluster N2 relative to State-1, although the tail region largely remains similar to that in State-1 (Fig. 3c). These conformational changes result in an M-distance of 16 Å (Fig. 3b) and an E-distance of 40 Å (Fig. 3d). State-3 distinguishes itself by housing a Q10 molecule with a headgroup that aligns more parallelly with the membrane and a contorted midsection of the Q tail. Although the M-distance remains consistent with that in State-2, the E-distance is reduced by 4 Å (Fig. 3d), a change attributable to its silkworm-like undulatory motion (Fig. 3g). In State-4, a Q10 molecule only partially fills the Q-channel (E-distance = 23 Å). Its headgroup aligns flush with the membrane surface (M-distance = 0), while the tail spans the entire lipid bilayer (Fig. 3b). Furthermore, the headgroup in State-4 is surrounded by a cluster of acidic, basic, and polar residues inside the Q-channel (Fig. 3f). These residues appear to create a highly polarized environment connected to the aqueous matrix, facilitating Q protonation. Taken together, our findings support a model where Q10 progresses through the Q-channel via peristaltic motion (Fig. 3g).

**Fig. 3.**
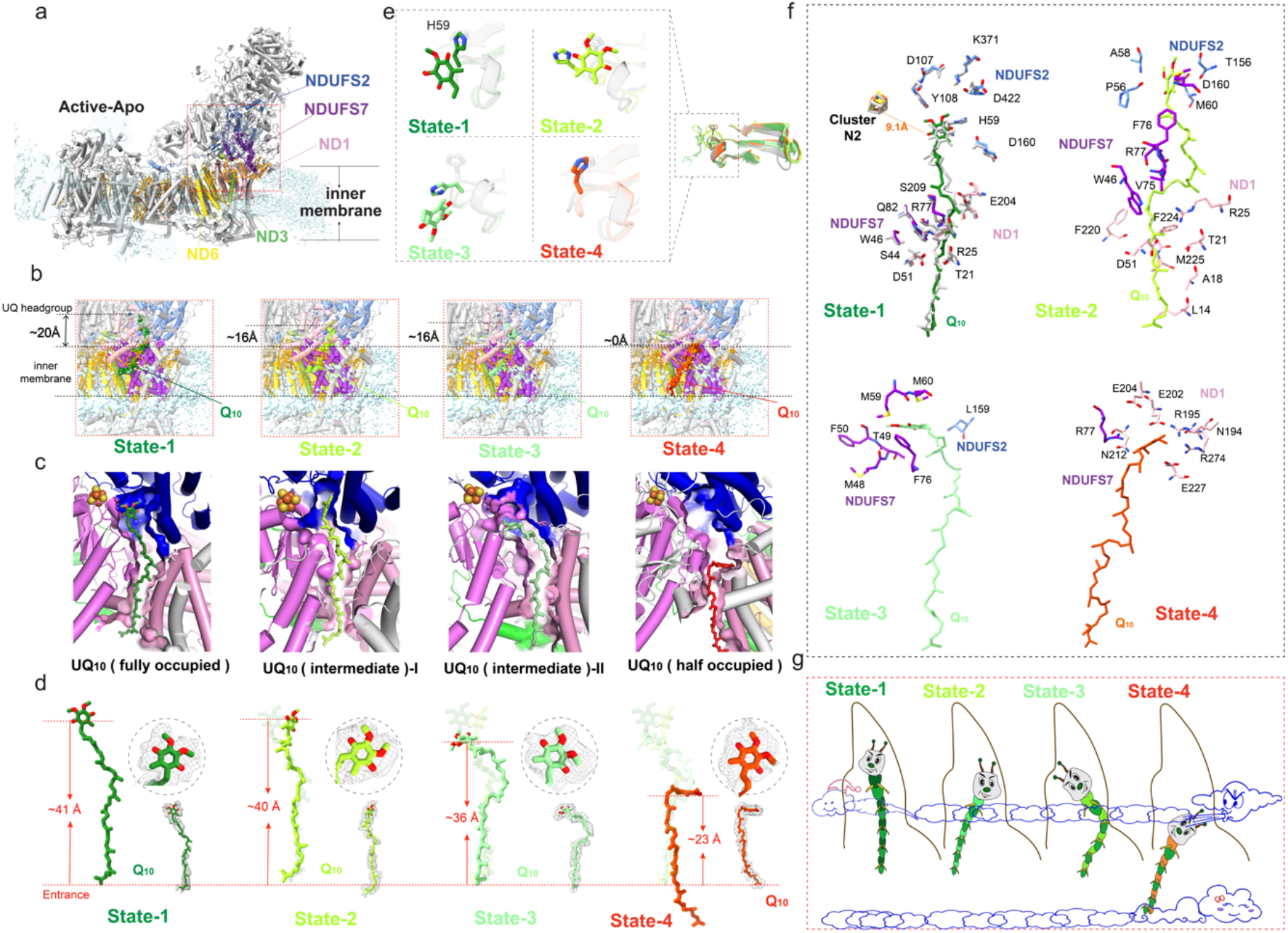
| High-resolution *in-situ* structures reveal multiple Q/QH_2_ binding states within the Q-binding channel. **a**, Cartoon models of CI active-apo state, with subunits constituting the Q-binding channel highlighted. **b,** Variations in distances between the Q_10_ headgroup and membrane surfaces across fully occupied, two intermediate, and half-occupied states. **c,** Spatial positioning of Q_10_ within the Q-binding channel for the four distinct binding states. **d,** Comparisons of three other binding states with the fully occupied Q_10_ (transparent stick). The distance from the Q_10_ headgroup to the quinone-binding channel entrance varies among different binding states. **e,** Superpositions of the four states with the active-apo state (grey) reveal not only conformational alterations in the Q_10_ headgroup and H59^NDUFS2^ (left panel) but also significant changes in long-range structures away from Q_10_ headgroup (right panel). **f,** Superimposition of the porcine *in-situ* supercomplex structure with the bovine CI (grey) incorporated into MSP nanodisks indicates that our fully occupied state resembles the active form of bovine CI with a Q_10_ fully occupying the Q-site (State-1). Residues interacting with Q_10_ in the other three binding states (State-2,-3 and -4). **g,** Schematic depiction of the silkworm-like undulatory motion of Q_10_ within the Q-binding channel.

### Active-Deactive Transitions in Complex I Under Different Conditions

Under physiological conditions and without substrates, mammalian CI transitions from an active, ready-to-catalyze state to a substantially deactive resting state^21,25,26^. During ischemia, a condition characterized by limited oxygen supply, the deactive state emerges due to the cessation of the electron transport chain^21^. To investigate the impact of varying ischemia levels on CI’s atomic-level structure in native mitochondria, we exposed porcine hearts to room temperature for durations of 0 minute (’fresh’), 40 minutes (’mild’), and over 4 hours (’harsh’) before mitochondrial isolation (Fig. 1b).

The active and deactive states of CI are delineated by distinct structural hallmarks, such as domain movements and conformational alterations around the Q-binding site and in proximal membrane-domain subunits^21^. Utilizing focused 3D classification, we analyzed these hallmarks from the supercomplex structures determined from the mitochondria under the three conditions (Extended Data Fig. 12 and 13). In the fresh sample, approximately 75% of CI in supercomplexes adopts an active state, whereas only ∼30% and ∼18% maintain this state under ’mild’ and ’harsh’ conditions, respectively.

Previous *in-vitro* structural studies suggest that the hallmarks of CI are collectively categorized into either the active or deactive form^23^. To further investigate whether there exist intermediate states involved in the transition from active to deactive, we performed additional focused 3D classification targeting the regions around the Q-site, resulting in 7 major intermediate classes. We built the models in these states and compared them to the canonical active and deactive structures (Extended Data Fig. 13 and Supplementary Table 1). The comparison unveils two distinct classes (class-0 and -7 in Supplementary Table 1) that fully conform to the hallmarks of the active and deactive states, respectively. By contrast, other classes exhibit only a subset of the hallmarks associated with the deactive state, suggesting that there exist various stages in the transition between the two extreme states in the native mitochondrial environment.

### Mechanistic Correlation of Rieske Domain Movement and Ubquinone Binding States in Complex III

CIII transfers electrons from ubiquinol to cytochrome *c* (cyt *c*) and contributes to the proton gradient for ATP synthesis^27,28^. Comprising three core subunits (cyt *b*, cyt *c1*, and the Rieske iron-sulfur protein (ISP)) along with eight auxiliary subunits, each CIII hosts four metal centers (hemes *bH* and *bL* in cyt *b*, the [2Fe-2S] cluster in ISP, and heme *c1* in cyt *c1*) and two Q-sites: Qi for Q reduction and Qo for QH2 oxidation^29–31^. From our high-resolution maps, with local resolutions between 1.8 and 2.4 Å (Extended Data Fig. 3 and 8) except for the dynamic ISP, we unambiguously identify all reactive centers and build atomic models of all endogenous ligands (Fig. 4a-d and Extended Data Fig. 9). Focused 3D classification reveals that endogenous Q10 ligands bound to both Q-sites adopt multiple conformations representing different reaction stages. Additional well-defined densities in the Q-binding pockets are identified as the structured lipids with their phosphate headgroups tightly bound to the surrounding protein regions. Tails of these lipids create a hydrophobic, dynamic environment for Q-binding and releasing.

**Fig. 4.**
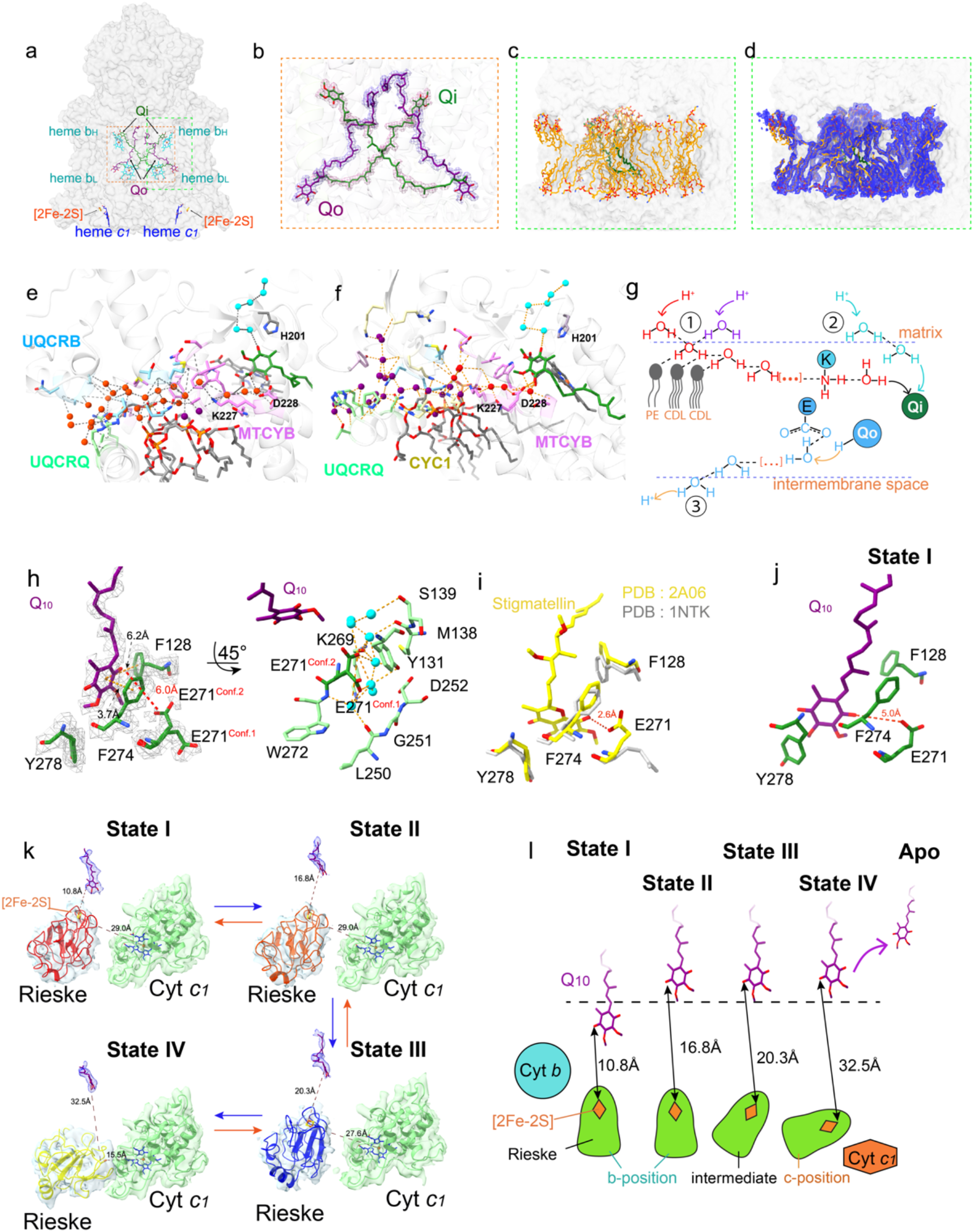
| Dynamic intermediates of CIII revealed by high-resolution *in-situ* cryo-EM. **a, b**, The structures resolve all reactive centers and endogenous Q_10_. **c,d,** The lipid molecules in the CIII Q-binding pocket and around the pocket entrance. These lipids are organized in a relatively ordered manner. **e,f,** Different views of the hydrogen-bonded network and the water chains for proton transfer. Red and purple spheres denote waters in the two branches of the bifurcated proton-influx path, while cyan spheres indicate waters in the single-wired proton-influx path. **g,** Schematic illustration of ① bifurcated proton-influx path ②, single-wired proton-influx path, and ③ proton-outflow path. **h,** Structural details of the Q_o_ binding site in the most abundant class. E271^MTCYB^ displays dual conformations in this class. Q_10_ and amino acids are colored in purple and green respectively. Blue spheres denote waters in the proton-outflow path near the Q_o_ site. **i,** Comparison of E271 conformations between the apo-form CIII (PDB:1NTK) and its complex with the Q_o_ inhibitor stigmatellin (PDB:2A06) from the same view showed in (**h**). **j,** Structural details of the Q-binding site in State-I with the closest distance to [2Fe-2S]. **k,** Four main Q-bound states of CIII as revealed by high-resolution *in-situ* structures. The Rieske sequentially moves from b-position to c-position to shuttle electrons. This movement is coupled with the Q-binding state at the Q_o_ site. Density maps for Q_10_ headgroups are shown as blue mesh, while those for the Rieske domain and Cyt *c_1_* are represented as transparent surfaces. **l,** Schematic representation for the coupling of ISP movement to Q-binding states.

During the Q-cycle, Q at the Qi site undergoes a two-step reduction, acquiring two protons from the mitochondrial matrix to form QH2^32^. Our structure uncovers a hydrogen-bond network near the Qi site, comprised of water molecules (Fig. 4e,f, Extended Data Fig. 14 and Supplementary Video 5), polar amino acid residues, and the phosphate heads of three lipids (two CDL and one PE) (Fig. 4g), which form a Grotthuss-competent system for proton transfer^20,33^. The network is divided into two hydrogen-bond chains primarily constituted of water molecules that link Q10 to the matrix (Supplementary Video 5). One of the two chains further branches above the three lipids that is directly involved in facilitating the proton transfer through the hydrogen network. This bifurcated chain engages two charged amino acids, K227 ^MTCYB^ and D228 ^MTCYB^, with D228 ^MTCYB^ linked to the 1-carbonyl of Q10 (Fig. 4e, f). The second chain, comprising 5 waters that surround H201^MTCYB^ (Fig. 4e, f), is directly connected to 4-carbonyl of Q10. These chains provide an explanation for how the rapid proton transfer process is achieved in the Qi site.

At the Qo site, QH2 releases two protons into the intermembrane space as part of its reaction cycle^34^. Our high-resolution map clearly resolves a prevalent Q-binding conformation at this site, characterized by π-π stacking interactions with F274^MTCYB^ (Fig. 4h). Similar to the Qi site, there also exists a hydrogen-bonded water chain near the Qo site for proton transfer. It is worth noting that the residue E271^MTCYB^, engaged in this water chain, exhibits clear dual conformations (Conf. 1 and 2) and likely plays a critical role in proton transfer (Fig. 4h). The two conformations resemble those in two previously reported crystal structures in the apo state^35^ (Conf. 1) and with the Qo-inhibitor stigmatellin bound^36^ (Conf. 2), respectively (Fig. 4i). Further structural comparison indicates that the headgroup of stigmatellin lies closer to the ISP than the endogenous QH2 captured in our *in-situ* structure. Focused 3D classification uncovers a less-populated (∼13%) Q-binding state (Fig. 4j) at the Qo site with its headgroup shifted ∼6 Å closer toward the ISP, compared to the dominant form (Fig. 4k, l). We propose that this state, rather than the dominant class, represents a transient electron-hopping phase from QH2 to [2Fe-2S] in ISP.

The transfer of electrons from QH2 at the Qo site to cyt *c1* is mediated by the movement of the Rieske domain^28,32^. While early structural studies hinted at its dynamic role in electron transfer, questions remain about how this motion is initiated, regulated and coupled with Q-binding states during the catalytic cycle. In addition to the apo form, we have resolved four distinct Q-bound states (Fig. 4k, l). In State-I, the Q10 headgroup is situated near the [2Fe-2S] cluster at a distance of 10.8 Å, with the Rieske domain exclusively adopting the b-position. In States-II, III, and IV, Q10 takes on the prevalent conformation revealed in our cryo-EM maps, while the Rieske domain undergoes a stepwise movement from the b-position towards the c-position, eventually making contact with cyt *c1* for electron transfer in State-IV. To further analyze the detailed conformational changes among these states, we measure two characteristic distances, one from [2Fe-2S] to Q10 and the other to heme *c1* (Fig. 4k). Our results suggest a possible sequential progression from State-I to State-IV, characterized by increasing [2Fe-2S]-Q10 distances and decreasing [2Fe-2S]-heme *c1* distances. Based on these observations, we propose a model that outlines the coupling between ISP movement and Q-binding states.

### Coupled Functional States of CI and CIII Due to Q Reoxidation States

To investigate the potential correlation between the conformational states of CI and CIII within the same supercomplex, we performed focused conformational analyses on paired CIII and CI from differently treated samples (Fig. 1b). Using the ‘fresh’ sample, CI predominantly exists in its active state, which is expected to yield a higher concentration of QH2 (Fig. 5b). Concurrently, we observed that the conformation of the ISP domain of CIII predominantly adopted the b-position. In the ’harsh’ sample, most CI particles are classified into the deactive form, and the ISP domain of CIII preferentially adopts the c-position (Fig. 5c). Interestingly, although CI from the ‘mild’ sample is predominantly deactive, the ISP domain of CIII largely remains in the b-position, maintaining its catalytic functionality for a short period even after CI becomes deactive (Fig. 5c). We attribute this observed behavior to a temporal delay in QH2 depletion, coupled with the contributory role of Complex II in partially replenishing the QH2 pool. However, if CI remains in its deactive state for an extended period, this could eventually lead to QH2 depletion, which may in turn induce a conformational change causing CIII’s ISP domain to predominantly adopt the c-position.

**Fig. 5.**
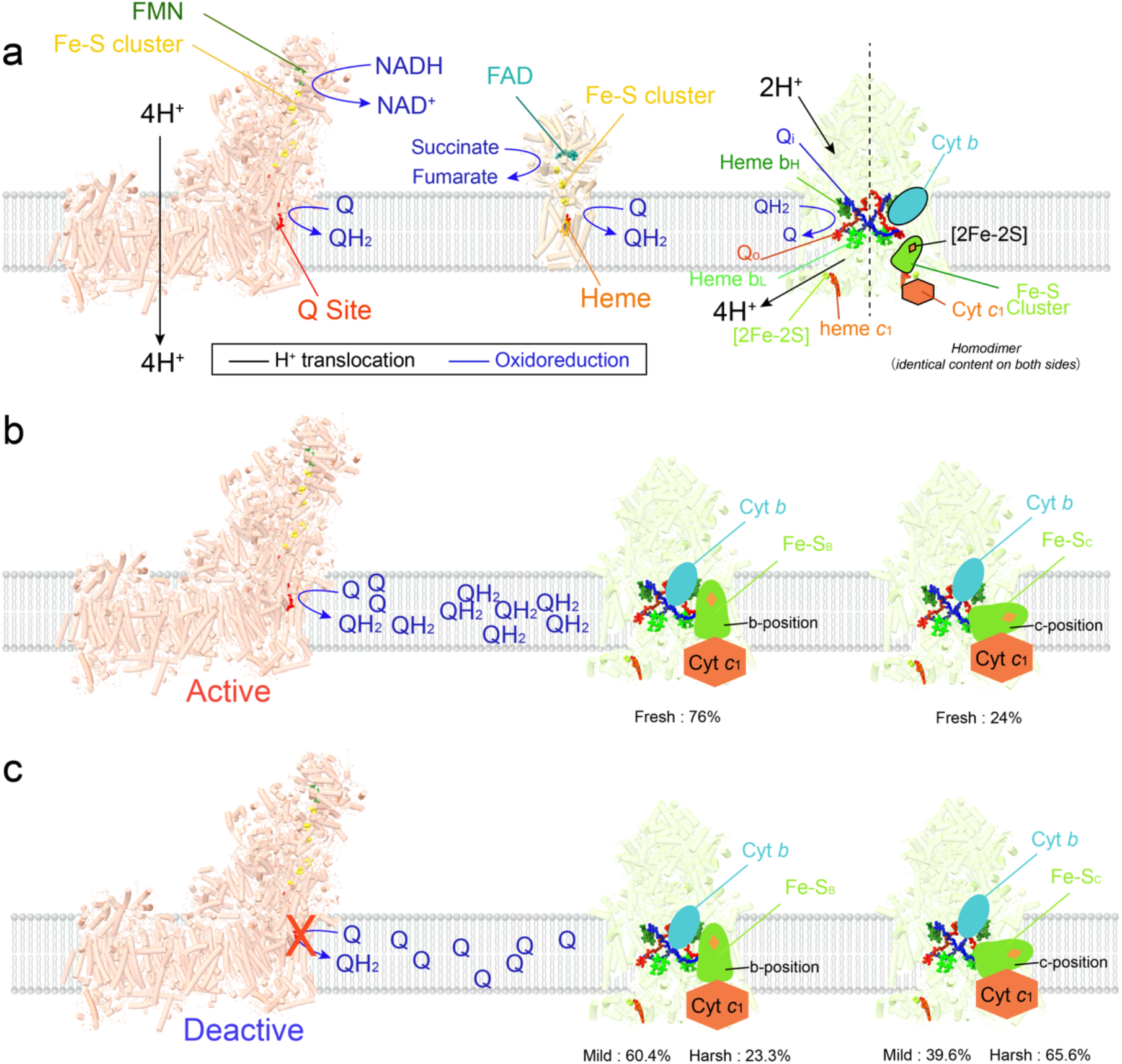
| Coupled functional states of CI and CIII due to Q reoxidation states. **a**, CIII utilizes QH₂ from either CI or II as its substrate. **b,** When CI is active, it utilizes NADH and Q to generates QH₂. Our structural analyses indicate that under these conditions, the Rieske domain of CIII predominantly adopts the b-position. **c,** In the deactive state of CI, QH₂ production ceases.

Prior to QH₂ depletion, the Rieske domain maintains its dominant b-position (including intermediate states similar to the b-position) for a period, and transitions to the c-position once QH₂ is fully depleted.

## Discussion

Our study employs *in-situ* cryo-EM imaging techniques to directly visualize supercomplex structures within mitochondria. This approach bypasses the limitations associated with traditional *in-vitro* purification methods, which could result in the loss of native ligands and physiological states, leading to compositional and conformational artifacts. Our approach preserves both the native membrane environment and the electrochemical proton gradient. This enables concurrent analysis of supercomplex structures in reaction at both atomic details and large-scale organizations, achieving a local resolution of sub-2 Å. Optimized *in-silico* classification of the *in-situ* cryo-EM data enables us to capture a series of dynamic ligand-binding states within the respiratory chain, eliminating the requirement for analogs or reaction inhibitors commonly used in conventional methods, and thereby providing new insights into the mechanisms of mitochondrial supercomplexes that reflect the physiological behavior. Intriguingly, the preservation of supercomplex structures in native membranes enables direct visualization of numerous surrounding lipids, which serve pivotal roles at the interaction interfaces among the complexes as well as within the Q-binding pockets. Encouragingly, these findings hold the potential for extended applications to elucidate the effects of many analogs, drugs and inhibitors on mammalian mitochondria, thereby shedding light on supercomplex behavior under various physiological and pathological conditions. Furthermore, our *in-situ* approach opens the door to the study of native structures of many membrane protein complexes at the organelle level.

## Acknowledgments

All cryo-EM datasets were collected at the Yale Cryo-EM Resource facilities (Science Hill and West Campus) and Brookhaven National Laboratory. We thank S. Wu, J. Lin and K. Zhou at Yale University and Liguo Wang, Jake Kaminsky, Guobin Hu at the Laboratory for BioMolecular Structure (LBMS) for their technical support in electron microscopy, as well as M. Gu, B. Ma, and Z. Huang for their assistance in preparing materials necessary for the manuscript. This work was supported by startup funds from Yale University and by grants from the National Institutes of Health, R35GM142959 awarded to K. Zhang, and S10OD023603 awarded to F. Sigworth at Yale University.

## Author contributions

K.Z. and J.Z. designed the project. W.Z. prepared all samples for cryo-EM/ET studies. W.Z. and K.Z. collected all cryo-EM/ET data. All authors contributed to cryo-EM/ET data processing, built the atomic models, analyzed the results, and prepared the manuscript.

## Materials and Methods

### Preparation of porcine mitochondria and cryo-EM grids

Mitochondria were isolated from porcine hearts following a modified protocol originally described by A.L. Smith^37^. Prior to mitochondrial extraction, the pig hearts were subjected to three distinct treatment conditions: (1) ‘Fresh’—immediately placed on ice for all subsequent procedures; (2) ‘Mild’—incubated at room temperature for 40 minutes and put on ice to quickly cool down for isolation; and (3) ‘Harsh’—incubated at room temperature for over 4 hours before cooling on ice. The isolated mitochondria were then resuspended in a solution containing 0.25 M sucrose, 10 mM Tris-buffered with H_2_SO_4_, 0.2 mM EDTA at pH 7.8. The suspension was adjusted to achieve a final optical density at 600 nm (OD_600_) of 1.3 AU.

For cryo-EM grid preparation, 3.3 μL of the mitochondrial suspension was applied to each Quantifoil hole carbon grid (R2/1, 300 mesh gold). Grids were incubated for 5 seconds within a Vitrobot Mark IV (ThermoFisher Scientific) chamber maintained at 8°C and 95% relative humidity. Excess solution was blotted using standard Vitrobot filter paper before the grids were rapidly plunged into liquid ethane at a temperature of approximately -170°C.

### Cryo-ET Data Collection

Grids were initially screened for optimal ice conditions using a 200-kV Glacios microscope (Thermo Fisher Scientific) at the Yale Science Hill Electron Microscopy Facility. Selected grids were subsequently transferred to a 300-kV Titan Krios microscope (Thermo Fisher Scientific), equipped with a Bioquantum Energy Filter and a K3 direct electron detector (Gatan), for high-resolution data acquisition at the Yale West Campus Electron Microscopy Facility. Automated data collection was facilitated using SerialEM software^38^. All images were captured in super-resolution mode, with a physical pixel size of 6.1 Å (effectively 3.05 Å in super-resolution). A total of eight tilt series were collected, targeting a relatively high defocus range, from -6 µm to -10 µm, for better contrast to guarantee a more reliable initial reconstruction. A grouped dose-symmetric scheme, spanning from - 60° to 60° at 2° increments, was employed for tilt series acquisition, with an accumulated dose of 100 e⁻/Å².

### Cryo-ET reconstruction and sub-tomogram averaging

Tomogram reconstruction was streamlined using custom scripts. Initial frame alignment was performed using MotionCorr2^39^, followed by micrograph binning at a factor of two. Tilt series stacks were generated using in-house scripts. All tilt series were aligned and reconstructed using AreTomo 1.2.5_40_. Initial CTF parameters were estimated via GCTF^41^ and cryoSPARC^42^. Raw micrographs and reconstructed results were visualized and diagnosed using IMOD^43^ and ChimeraX^44^.

Individual supercomplex particles were picked in EMAN_2 45_. Metadata preparation yielded 12,000 sub-tomogram particles in RELION_4 46_ with a binning factor of 2 (pixel size 12.2 Å). Following two rounds of 3D classification, 806 supercomplex particles were selected for final refinement, resulting in a 37 Å sub-tomogram averaging map. Resolution was assessed using Fourier Shell Correlation (FSC) with a threshold of 0.143 in RELION4^46^. The averaged map was backprojected onto the original tomogram using the *subtomo2Chimera* code, available at https://github.com/builab/subtomo2Chimera. A summary of acquisition parameters and particle numbers is provided in Supplementary Information Table 1.

### Single-particle cryo-EM data collection

Automated data acquisition was executed on either a Glacios or Titan Krios electron microscope (Thermo Fisher Scientific). The Glacios was equipped with a K3 direct-electron detector (Gatan) and operated at 200 kV at a pixel size of 0.434 Å in super-resolution mode, with an objective aperture of 100 μm employed. The Titan Krios, also equipped with a K3 direct-electron detector, was operated at 300 kV at a pixel size of 0.416 Å in super-resolution mode with a Gatan energy filter. Automatic data collection was facilitated using the SerialEM software package^38^. Multi-shot acquisition parameters were set at 3×3 holes per imaging location, with four exposures per hole at 200 kV and five exposures per hole at 300 kV. The total electron dose was fractionated to 42 e⁻/Å² for the Glacios and 50 e⁻/Å² for the Titan Krios, distributed across 45 frames at 40 ms per frame. Defocus parameters ranged from -1.0 μm to -3.0 μm for the 200 kV dataset and from -1.3 μm to -3.0 μm for the 300 kV datasets. Details of the data collection are summarized in Supplementary Information Table 2-7.

### Pre-processing

For all datasets, motion correction was performed by MotionCor2^39^ or cryoSPARC^42^. The contrast transfer function (CTF) of each motion-corrected micrograph was estimated using Gctf^41^ or cryoSPARC^42^. Particles were picked by Gautomatch or cryoSPARC using an iterative sorting strategy as described below. Cryo-EM scripts used for real-time data transfer and on-the-fly pre-processing can be downloaded at https://github.com/JackZhang-Lab.

### Overall particle selection and sorting strategy

Due to the challenges posed by low signal-to-noise ratios (SNR) and a highly congested macromolecular environment (Extended Data Fig. 2a), traditional particle selection methodologies were insufficient for generating datasets amenable to reliable two-dimensional (2D) classification, *ab-initio* three-dimensional (3D) reconstruction, and subsequent local refinement. To address this issue, we implemented an iterative strategy to optimize particle selection. The approach involved multiple rounds of iterative 2D particle picking, 2D classification and 3D analyses including ab-initio 3D reconstruction, 3D classification and multi-level local refinement. Different from the conventional particle selection approach, our new strategy used Gautomatch and cryoSPARC^42^ for template matching to gradually increase the resolution of 3D projections as the reconstructions were progressively improved over cycles. We use various sources of references and cross-validate of the final results among them. To maximize the yield of high-quality particles, particles from the classes that show clear features of supercomplexes in all cycles were merged for subsequent 3D cross-classification. More details of the strategy are explained in the following sections.

### Initial 3D reconstruction with surrounding membranes (type-A)

Conventional 2Dclassification failed to generate meaningful class averages using images selected from our *in-situ* cryo-EM micrographs of mitochondria for three main reasons: (a) thick samples that lead to low SNR ratios and large defocus variations, (b) crowded environment that affects particle detection and alignment, and (c) strong membrane signals that dominate the alignment, leading to blurred averages of protein regions (Extended Data Fig. 2b).

To address this, we initially utilized the strong membrane signals and focused on the side views surrounded by membranes using 2D classification. These side views in principle contain sufficient orientational information for a complete 3D reconstruction. At the outset, protein signals were completely averaged out in the 2D classification, whereas the membranes were well aligned due to the strong side-view signals (Extended Data Fig. 2c). We then conducted multiple cycles of 2D classifications to focus only on particles exhibiting clear membrane signals.

Through comprehensive 2D analyses, we found regions potentially harboring mitochondrial supercomplexes exhibit special features of local curvature. Specifically, these regions are characterized by membrane signals appearing to be concave towards the matrix direction, indicative of the presence of CIII (Extended Data Fig. 2c-e). By merging particles from classes with characteristic concave membranes surrounding CIII and conducting further 2D classification, we achieved improved 2D averages showing clear membrane features around CIII (Extended Data Fig. 2d). Notably, extra protein densities adjacent to CIII were obvious, likely accounting for CI or CIV densities. However, it had been unclear how many types of respiratory supercomplexes exist in native mitochondria, and whether CI, CIII and CIV always appear in the form of supercomplexes or just partially.

To further address these observations and obtain unbiased density maps, we employed four independent methods to generate initial references: (1) cryo-ET sub-volume averaging (Extended Data Fig. 1), (2) *ab-initio* reconstruction using particles assigned to the 2D averages with visible protein densities (Extended Data Fig. 2f) and characteristic CIII membrane features (Extended Data Fig. 2g), (3) *ab-initio* 3D reconstruction using particles after membrane subtraction (Extended Data Fig. 2h), and (4) models generated from random selection of unsorted particles or random noise (Extended Data Fig. 2i). All these references were combined for 3D classification and subsequently used for local refinement and focused classification (Extended Data Fig. 2i). Given that *in-situ* cryo-EM datasets are more heterogeneous than conventional single-particle datasets, we included ’false references’ generated from approach (4) for better classification. Finally, particles corresponding to classes showing clear features of type-A supercomplex were re-extracted and merged for additional classification and refinement (Extended Data Fig. 2i).

### Cross-classification of multiple supercomplexes

Around the reconstructed supercomplex I_1_III_2_IV_1_ (type-A) map, we observed extra densities, clearly suggesting more proteins bound to the type-A supercomplex to form larger supercomplexes. We suspected more types of supercomplexes exist in native mitochondrial membranes. Preliminary results from both large single-particle 3D classification at low resolutions and cryo-ET sub-volume averaging confirmed our speculation. To further improve the accuracy of 3D classification for high-resolution refinement, we deliberately provided references generated from random sub-datasets using discarded particles from previous cycles. These ’false references’ served to randomly ’absorb’ low-quality and falsely picked particles, leading to a relatively clean dataset for the target class. We then accumulated particles classified into good classes, defined by clear secondary structures, over multiple cycles. Due to the crowded mitochondrial environment, misclassified and misaligned particles were always present. To address this, we re-organized the particles by merging those that fell into classes generating similar 3D maps. We selected multiple references from different classes, including those considered ’bad,’ and re-performed 3D classification on each sub-dataset. Afterward, we re-combined a subset of different classes and re-classified them. Based on the results from multiple subsets after re-classification, we merged all the particles belonging to a specific target from previous cycles and performed an additional cycle of 3D classification on this merged dataset. This additional classification used high-resolution references generated from previous classification cycles and local refinement to discard low-quality or misclassified particles.

After numerous rounds of cross-classification followed by local refinement, we identified various other types of supercomplexes, including the other three major classes: type-B (I_1_III_2_IV_2_), type-O (I_2_III_2_IV_2_), and type-X (I_2_III_4_IV_2_). Other classes such as I_1_III_2_ and I_4_III_4_IV_4_ were not subject to further refinement in this study due to their low population. Subsequently, we cross-validated our classification results by providing a set of references lacking the correct form of the supercomplex for sub-classification on each class. We also performed additional reference-free 2D classification after reference-based 3D classification and refinement. This allowed us to visualize the distinct features of the four major classes from 2D averages directly, without imposing any references. Only datasets converging to the correct form of supercomplex, regardless of the initial references used, were included in the final multi-level local refinement and focused 3D classification.

### Multi-level local refinement and focused 3D classification

A hierarchical masking strategy was employed for local refinement on all four major types of supercomplexes. Specifically, the mask size was incrementally reduced to focus on distinct regions of each type of the respiratory supercomplexes, ensuring stable local refinement. We partitioned the type-A supercomplex into five principal domains: (a) Complex I hydrophilic region, (b) Complex I hydrophobic region, (c) Complex III, (d) Complex IV, and (e) lipid environment.

Before the multi-level local refinement, the type-A supercomplex was refined to 3.39 Å overall using images binned two times (1.664 Å/pixel after binning) with 1,113,902 high-quality particles. This also included type-B, type-O, and type-X, as they all share the type-A region. We re-centered and re-extracted these particles, generating 1,050,463 particles for subsequent local refinement (particles near the edges were excluded after re-extraction). Initially, the resolutions of CI, CIII, and CIV worsened slightly (∼3.5 Å) after the first cycle of refinement using the unbinned particles (0.832 Å/pixel). Further improvement was achieved by optimizing several local refinement parameters, including optimization of mask sizes, global CTF, local CTF refinement, local angular refinement, and non-uniform refinement^47^.

By iteratively applying these techniques, the maps of the hydrophilic region of CI, the hydrophobic region of CI, CIII, and CIV were refined to average resolutions of 2.46 Å, 2.58 Å, 2.31 Å, and 2.66 Å, respectively (Extended Data Fig. 3a-c, Supplementary Information Table 2-7). Even smaller regional masks, focused on CI and CIII, further improved local resolutions. Local resolutions in the majority of the protein regions of CIII ranged from 1.8 to 2.4 Å (Extended Data Fig. 3c). Focused classification and refinement on specific subdomains, like the ubiquinone/ubiquinol binding sites, yielded further improvements that aided in model building. For more complex regions, such as the lipid environment surrounding the transmembrane regions of the supercomplexes and ubiquinone/ubiquinol binding sites, additional levels of focused classification and local refinement were performed. To ensure seamless integration of adjacent regions, all masks were manually created so that pairs of adjacent masks contain sufficiently large areas for the generation of final composite maps using the smaller regions individually refined. All locally refined segments were integrated into a comprehensive composite map in ChimeraX^44^.

Similar multi-level refinement approaches were applied to determine the structures of other forms of respiratory supercomplexes. Detailed parameters and refinement results are summarized in Extended Data Fig. 3-6 and Supplementary Information Table 2-7.

### Membrane signal detection, modeling and suppression

One of the critical bottlenecks limiting high-resolution cryo-EM reconstruction of membrane proteins in their native environment is the severe signal interference from surrounding membranes. This interference can significantly affect multiple steps in cryo-EM data analysis, including *ab-initio* reconstruction, Euler angle determination, and 2D and 3D classification, as well as refinement of alignment parameters. To address this issue, we developed a computational toolkit to detect membrane signals from 2D averages, estimate the local geometry of detected membranes, and suppress or remove these signals to substantially improve the alignment reliability of mitochondrial complexes in native membrane environments.

Initially, we generated a series of 15-30 computationally simulated 2D projections of lipid bilayers, with local curvatures ranging from 0 nm^-1^ to 0.02 nm^-1^. These simulated 2D membranes served as templates for detecting the side-view signals of mitochondrial membranes using Gautomatch. Subsequently, 3-5 cycles of 2D classification were performed to discard low-quality and non-membrane particles, resulting in a subset of particles showing clear side views of lipid bilayers. We then estimated the approximate orientation and center of each individual lipid bilayer based on its corresponding 2D average. Local curvature was determined by maximizing the cross-correlation between each 2D average and a series of simulated lipid bilayers. These curves were rotated and translated using alignment parameters from 2D classification generated by cryoSPARC^42^. Centers of each membrane segment were refined by maximizing the normalized cross-correlation between the raw image and transformed 2D average. Using these estimated parameters, the principal signals of each membrane segment were approximated by locally averaging the image intensities along the membrane curve within a soft mask, which was ∼25% larger than the typical lipid bilayer we estimated. Membrane signals that had previously dominated the alignment were subtracted from the raw images to enhance protein signal contributions for subsequent reconstruction, alignment, classification, and local refinement. This improved the signal contributions from protein regions, akin to the critical effects observed in our previously described microtubule signal subtraction method^48,49^. Finally, alignment and classification parameters were applied to the raw images along with membrane signals for subsequent classification, focused classification and local refinement.

### Membrane modeling and geometry analysis

The *in-situ* mitochondrial respiratory chain complexes largely preserved the native state of the membrane architecture, as evidenced by exceptionally clear density maps (Extended Data Fig. 7). This high fidelity in density was observable in both the final 3D reconstructions and the post-3D-refinement 2D class averages, enabling direct modeling of native membrane structures.

The model building for the inner membrane structures surrounding the mitochondrial supercomplexes involves a four-step procedure. First, discrete points are sampled from the raw signals in a given density map—such as the type-A supercomplex—based on binarized membrane density. A 2D plane is fitted by least-square minimization; the normal vector of each supercomplex is estimated and the coordinate system is rotated so that this vector aligns with the Z-axis. Second, these sampled discrete points are utilized to generate two smooth, curved surfaces, each with a thickness of ∼4 nm. Third, planar phospholipid bilayer structures are generated to match the geometry of these estimated surfaces. Finally, the information from the second and third steps is integrated to geometrically deform each planar membrane structure into a smooth, curved surface.

To optimize the initial sampling for membrane model building, we categorized the membrane structures surrounding the protein into three distinct categories: structured lipids, surface-associated lipids, and generic bilayer lipids. The first category, “structured lipids,” includes lipids that are closely associated with the protein’s transmembrane regions. This close association enables the identification and direct atomic-level modeling of these specific lipid species, which have also been observed in previously reported structures purified using detergent. The second category, “surface-associated lipids,” comprises lipids situated around the immediate periphery of the protein, forming a pseudo-lattice structure. Within this lattice, partial phosphatidyl head groups and hydrophobic tails can be discerned. Our *in-situ* density maps allow us to unambiguously determine the location of individual lipids in this category; however, the current quality of the density maps does not permit the identification of the specific types of lipids present. The third category, “generic bilayer lipids,” represents a region farther from the protein where only the density features corresponding to the bilayers could be observed. A generic phospholipid membrane model is employed to approximate the likely horizontal positions of the phosphatidyl headgroups. Due to the fluid nature of the lipid bilayer and the high level of noise in the density maps, the central positions of these generic bilayer lipids may still vary among different sub-classes even after focused classification. However, the average geometric features and the central locations of the membranes are notably consistent across each of the four major types of supercomplexes. Therefore, these generic bilayer lipids are used solely for calibrating the central locations and orientations of the phospholipid bilayer, rather than representing the actual positions of individual phospholipid molecules within each supercomplex’s bilayer. This approach aids in analyzing the overall geometric changes among the supercomplexes, albeit not at the level of individual phospholipid molecule structures.

To achieve a sufficiently smooth model for the generic bilayer lipids, we performed real-space refinement of the initial structures using the Coot software^50^. The refined structures were subsequently subjected to further smoothing using a local Gaussian filter to minimize residual noise in localized membrane regions. This step enabled the precise estimation of the contour map and the local curvature at each point (Fig. 2e-h). We utilized the CHARM-GUI web service^51^ to generate a simulated rectangular planar phospholipid bilayer. This planar structure was then mapped onto the curved surfaces that were obtained after Gaussian smoothing. This mapping process yielded a curved membrane model that optimally fit the density map. From these estimated surfaces, information about the local geometry of the membranes surrounding the mitochondrial supercomplexes could be directly retrieved for subsequent geometry analyses and comparisons.

### Model building, refinement, and validation

The atomic models were built manually using Coot^52^. First, high resolution structures of bovine CI (PDB code: 7QSK), bovine CIII (PDB code: 2A06) and bovine CIV (PDB code : 5XDQ) were fitted into corresponding map as a rigid body using ChimeraX^44^. Then the fitted model was manually mutated, adjusted and real-space refined to correct errors in local regions to best match the density maps using Coot^52^. The final model was refined using phenix.real_space_refine^53^ with geometric constraints. Figures were generated using UCSF ChimeraX^44^ and Pymol (The PyMOL Molecular Graphics System, Schrödinger, LLC).

### Data availability

All cryo-EM maps and atomic coordinates for mitochondrial supercomplexes have been deposited in the Electron Microscopy Data Bank (EMDB) and the Protein Data Bank(PDB), including entire and locally refined maps of type-A supercomplexes under accession codes EMD-42231/PDB-8UGP, EMD-42225/PDB-8UGH, EMD-42226/PDB-8UGI; type-B supercomplexes under EMD-42227/PDB-8UGJ; type-O supercomplexes under EMD-42230/PDB-8UGN; type-X supercomplexes under EMD-42233/PDB-8UGR. Different classes of CI under EMD-42165/PDB-8UEO, EMD-42166/PDB-8UEP, EMD-42167/PDB-8UEQ, EMD-42168/PDB-8UER, EMD-42176/PDB-8UEZ, EMD-42169/PDB-8UES, EMD-42170/PDB-8UET, EMD-42171/PDB-8UEU, EMD-42172/PDB-8UEV, EMD-42173/PDB-8UEW, EMD-42174/PDB-8UEX, EMD-42175/PDB-8UEY. Different states of CIII under EMD-42221/PDB-8UGD, EMD-42222/PDB-8UGE, EMD-42223/PDB-8UGF, EMD-42224/PDB-8UGG, EMD-42229/PDB-8UGL. High resolution representative of complex under EMD-42143/PDB-8UD1, EMD-42228/PDB-8UGK, EMD-42229/PDB-8UGL.

### Code availability

All codes involved in general cryo-EM/ET data processing and structure determination of membrane proteins will be publicly available at: https://github.com/JackZhang-Lab.

---

**Extended Data Fig. 1.**
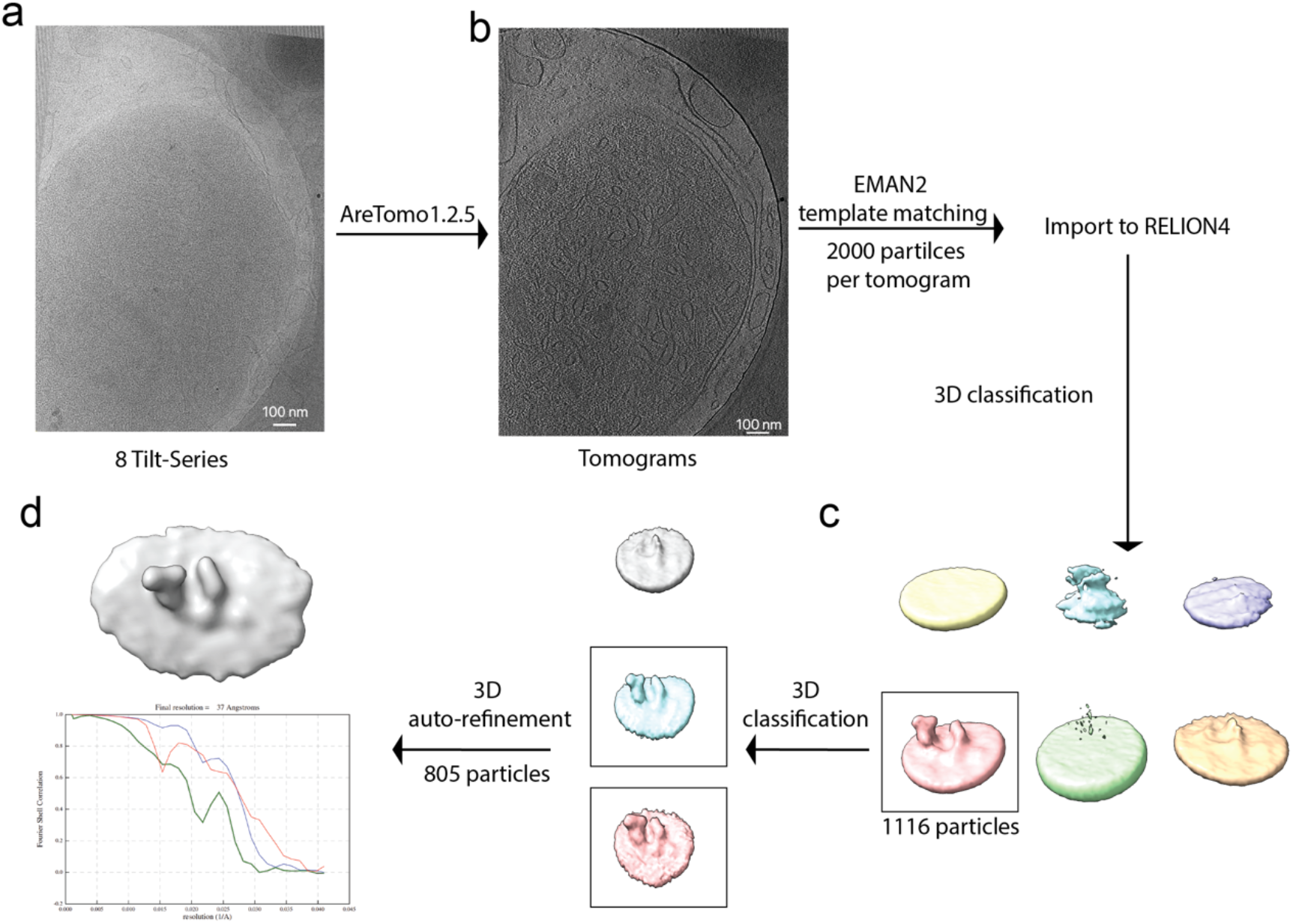
| Cryo-ET data processing flowchart. **a**, Representative zero tilt image from 8 tilt series. **b**, Representative slice of the reconstructed tomogram. **c**, Intermediate results of the sub-volume averaging by Relion 4.0. **d**, Final reconstructed map and FSC curve of the type-A supercomplex.

**Extended Data Fig. 2.**
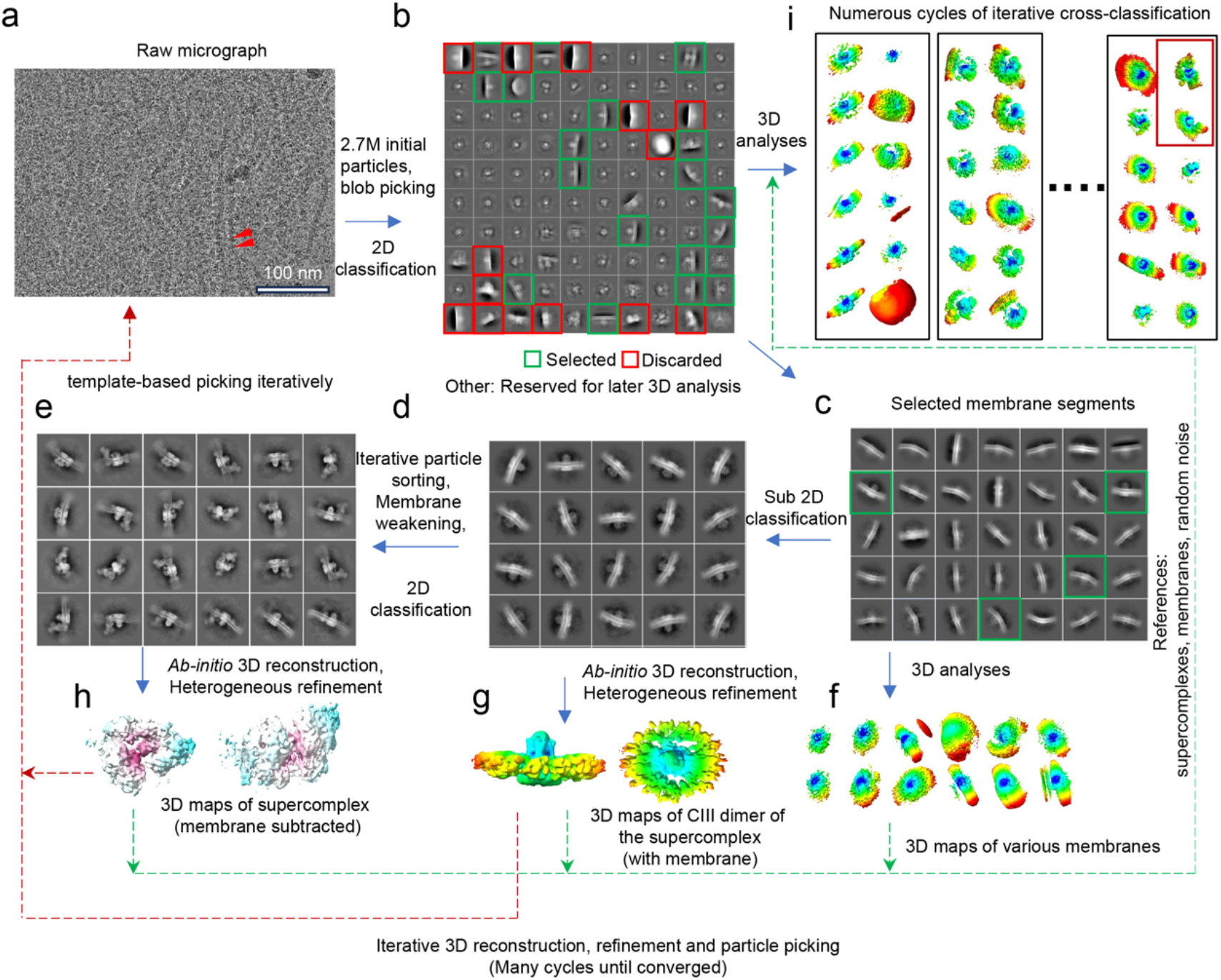
| Schematic Overview of Initial Single-Particle Cryo-EM Data Analysis Workflow. **a**, High-magnification micrograph exemplifying a typical single-particle cryo-EM sample. **b**, Preliminary results of reference-free 2D classification. **c**, Representative outcomes of 2D sub-classification focused on particles displaying discernible side views of mitochondrial membranes. **d**, Illustrative 2D classification of mitochondrial supercomplex particles derived from the prior 2D classes; features reveal distinct concave membrane morphologies in the CIII regions. **e**, Exemplary 2D class averages after extensive particle sorting and membrane signal weakening. **f-h**, 3D reconstructions generated using particles selected from steps (**c**), (**d**), and (**e**), respectively. **i**, Typical results of the iterative cross-3D classification. Reference maps and particle classification/alignment parameters were progressively refined over multiple cycles. Upon reaching this stage, reliable 3D reference models were obtained for subsequent multi-level refinement and focused classification. Particles classified into supercomplexes were merged for all ensuing data processing steps.

**Extended Data Fig. 3.**
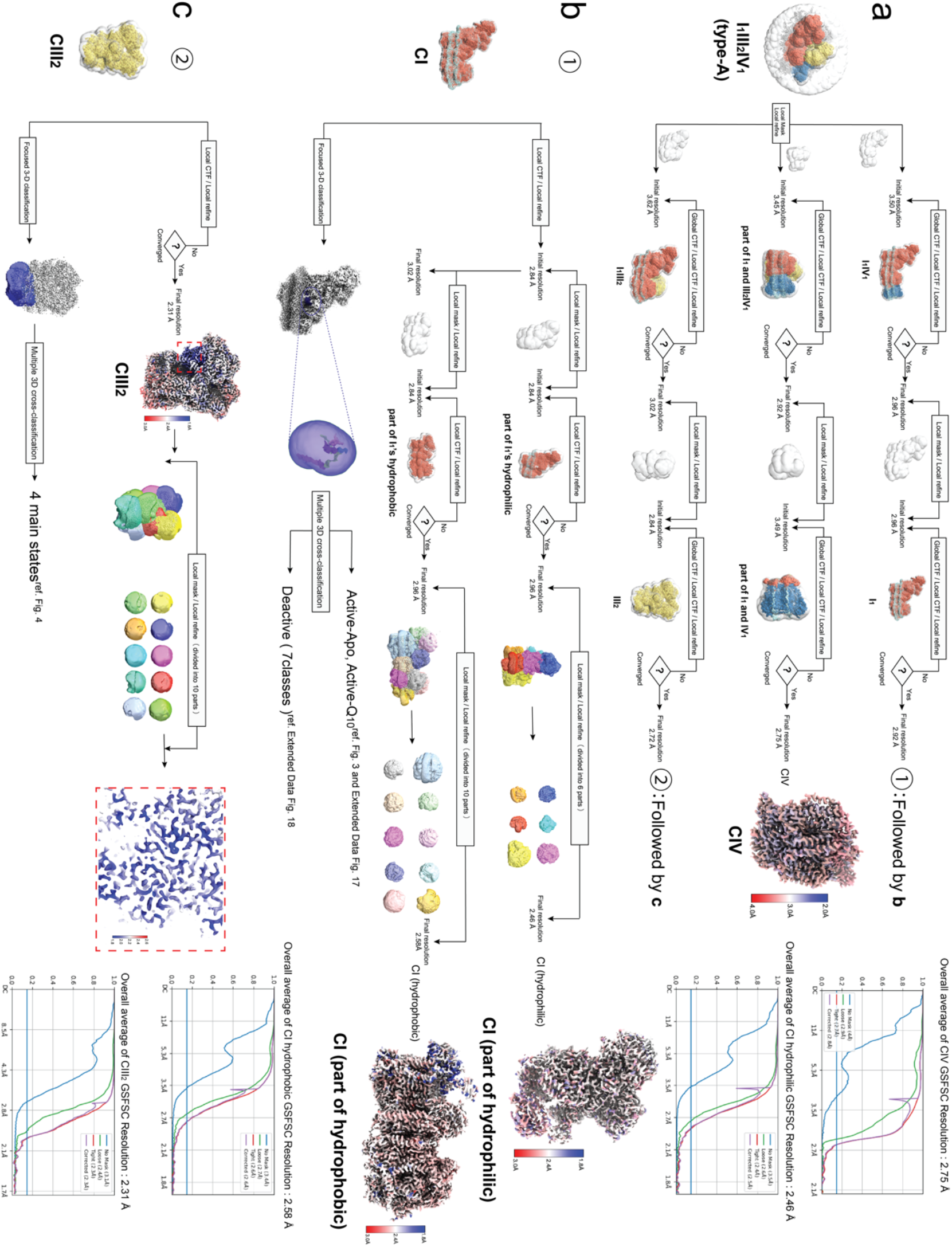
| Schematic of the workflow for multi-level refinement and focused classification of type-A supercomplex. **a**, Multiple cycles of local refinement that generated maps of CI, CIII, CIV. **b**, Multi-level local refinement of CI and focused classification of Q_10_ region. **c**, Multi-level local refinement of CIII and focused classification of the Rieske domain.

**Extended Data Fig. 4.**
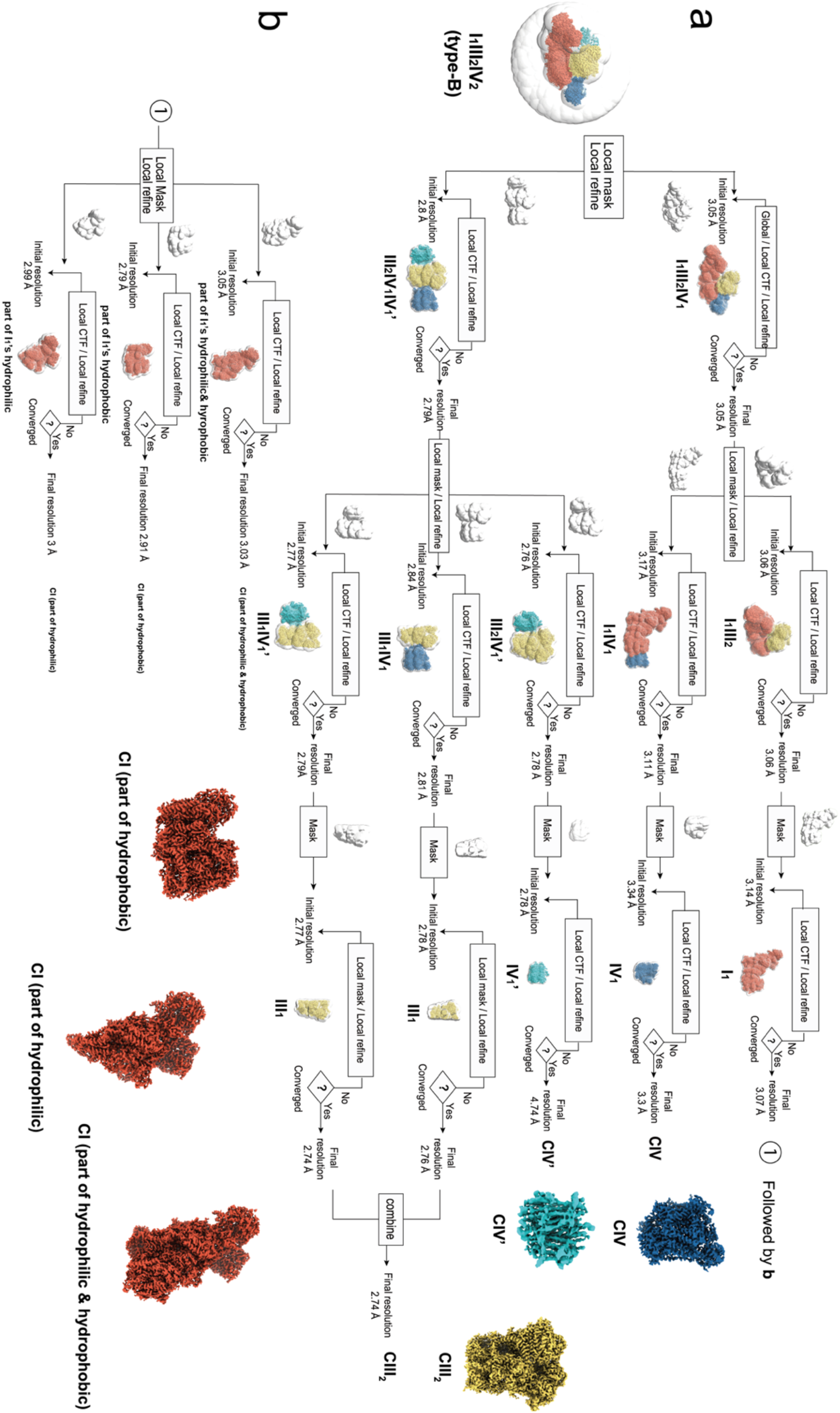
| Schematic of the workflow for multi-level refinement and focused classification of type-B supercomplex.

**Extended Data Fig. 5.**
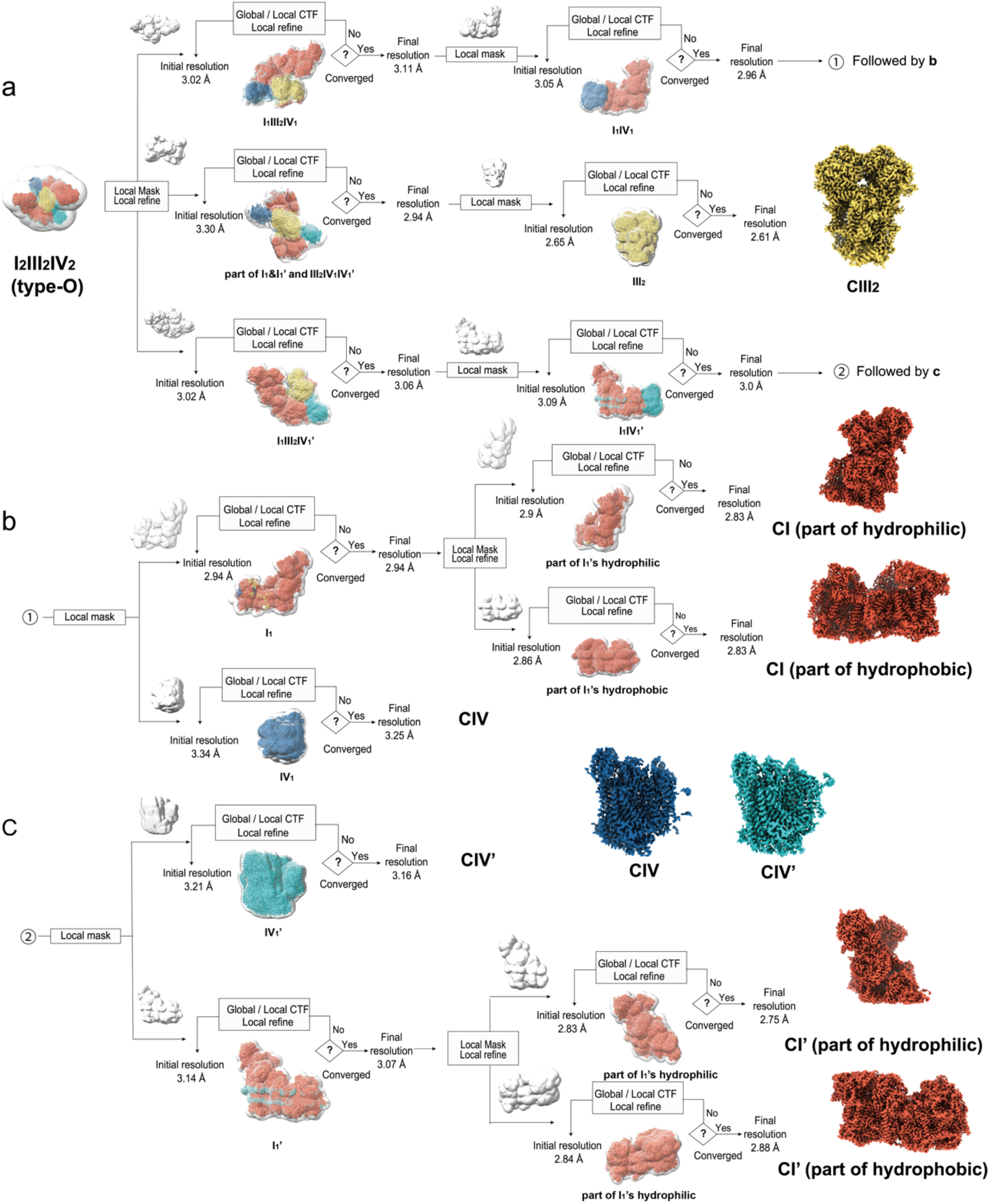
| Schematic of the workflow for multi-level refinement and focused classification of type-O supercomplex.

**Extended Data Fig. 6.**
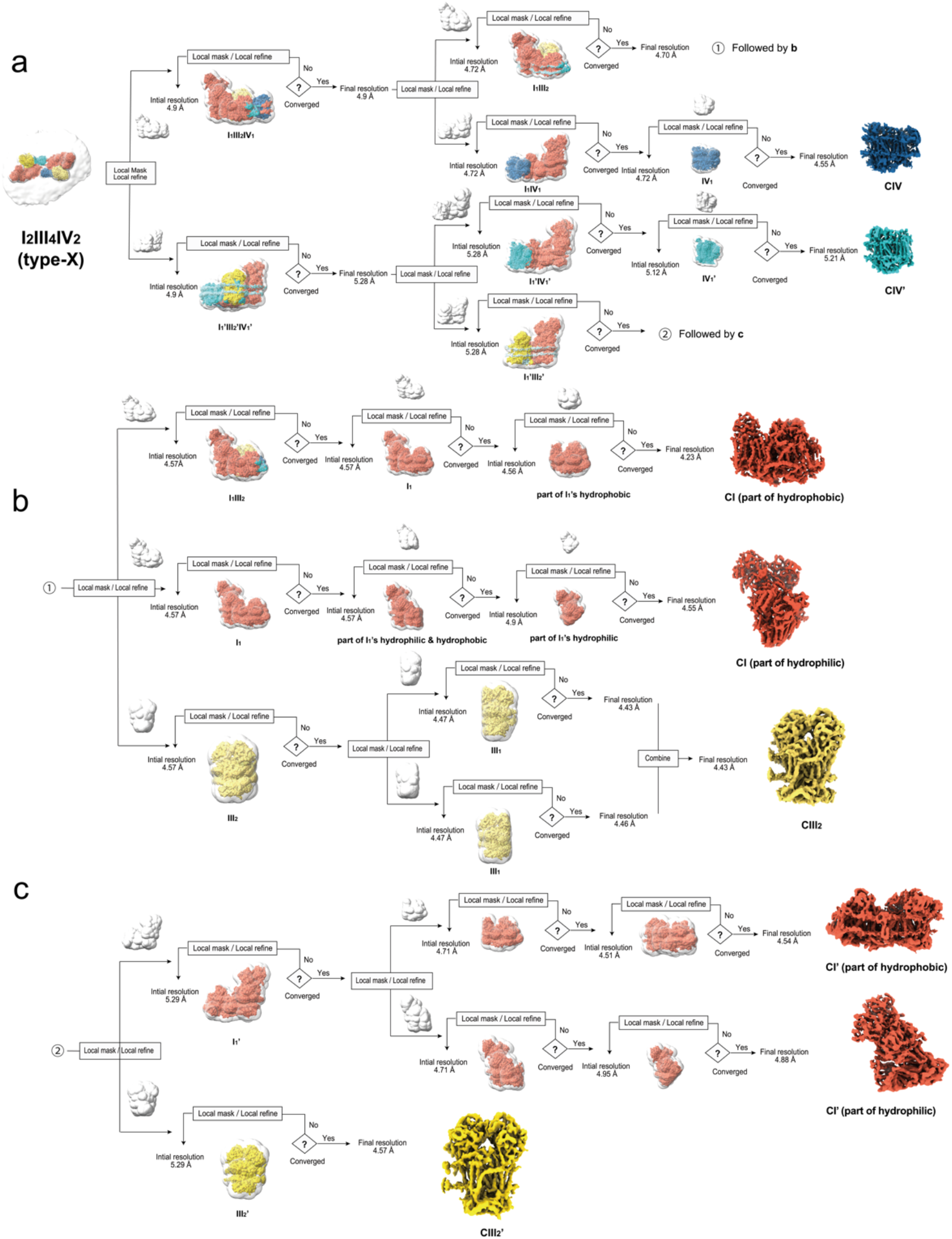
| Schematic of the workflow for multi-level refinement and focused classification of type-X supercomplex.

**Extended Data Fig. 7.**
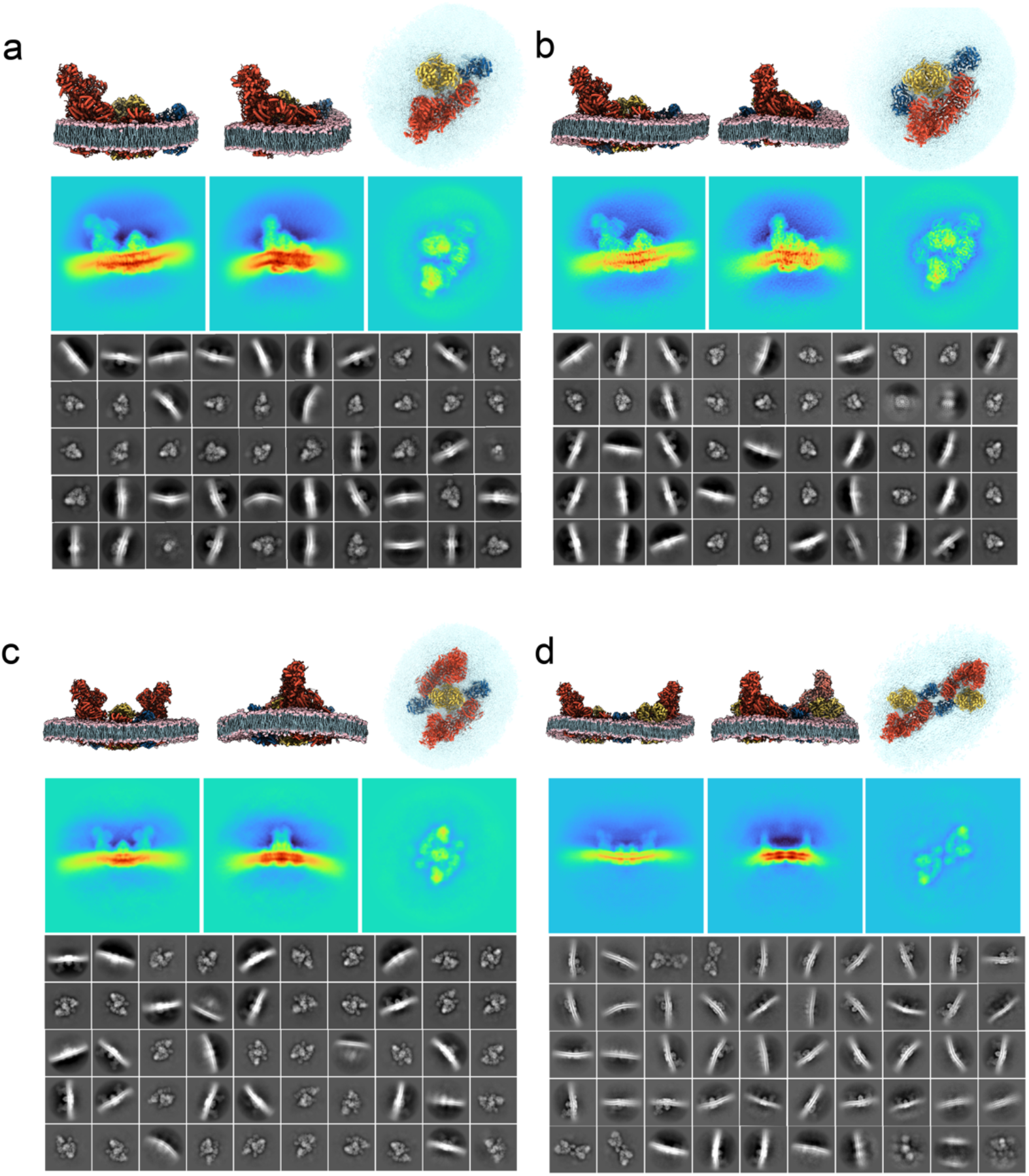
| Evaluation of the 3D classification by post-3D-refinement 2D classification. **a-d**, Atomic models (top) from three representative views, the corresponding projections from the same orientations (middle) and representative reference-free 2D class averages of type-A (a), -B (b), -O (c) and -X (d) supercomplexes after 3D classification and refinement.

**Extended Data Fig. 8.**
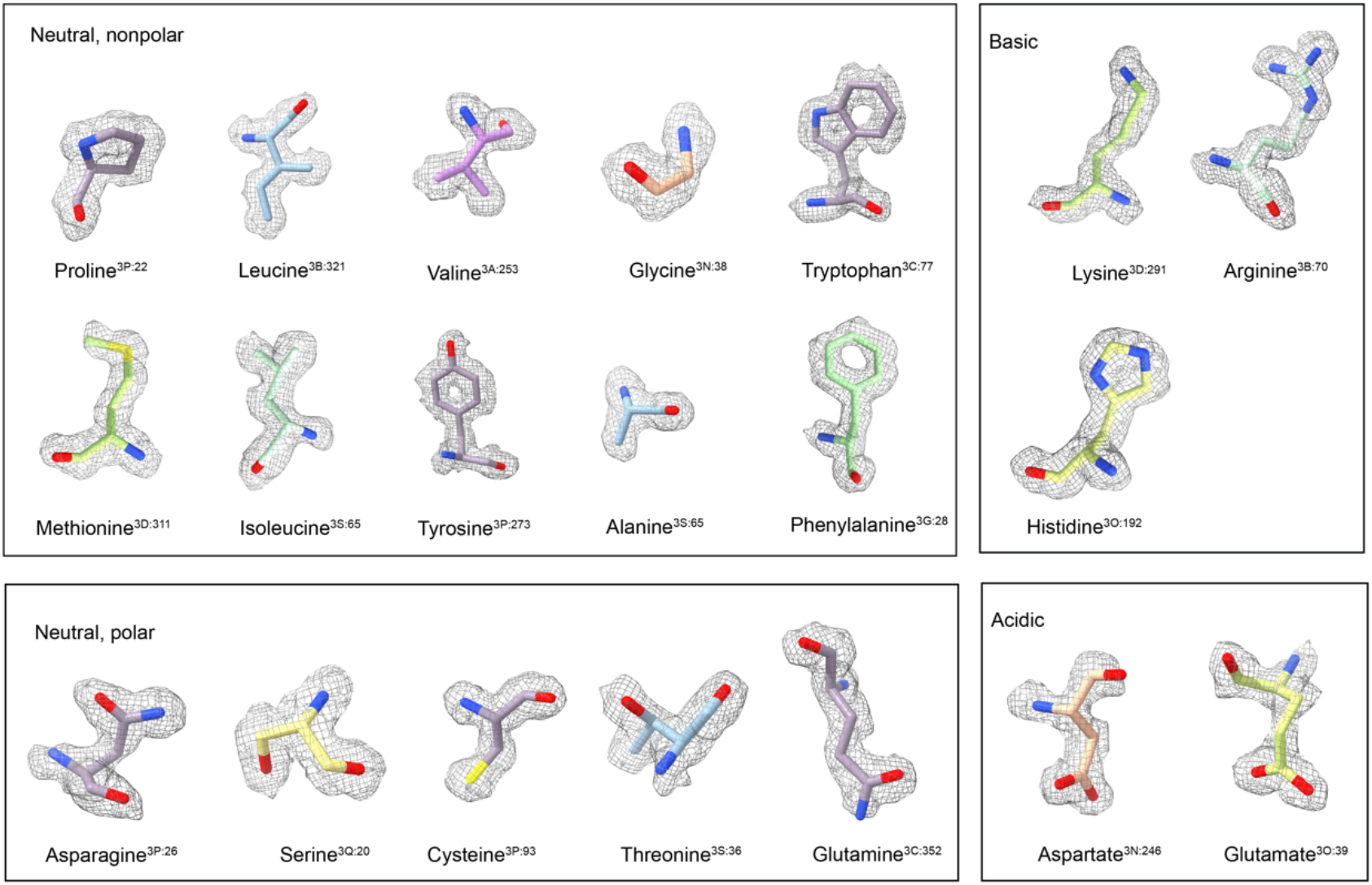
| Representative densities in CIII of all types of residues demonstrate high-resolution features enabling unambiguous rotamer determination. Notably, the distinct holes in the six-element rings of tryptophan, phenylalanine, and tyrosine are prominently visible.

**Extended Data Fig. 9.**
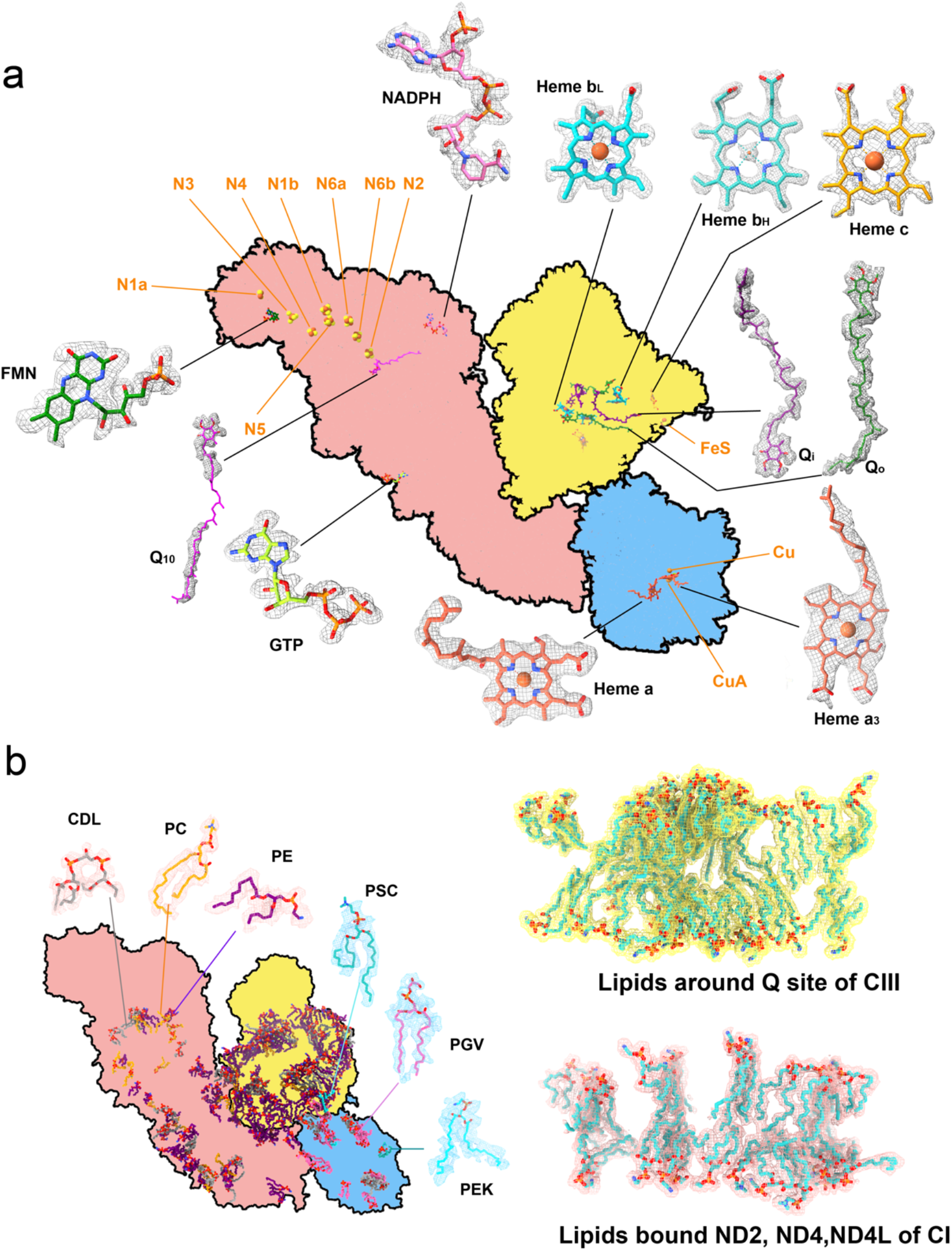
| Endogenous cofactors and representative structured lipids revealed by the high resolution *in-situ* cryo-EM structures. **a**, Representative density maps of endogenous cofactors and atomic models fitted. The cartoon in the center represents the contour of the type-A supercomplex projection with the ligand position assigned. **b**, Structured and associated lipids identified from the high-resolution cryo-EM density of type-A supercomplex.

**Extended Data Fig. 10.**
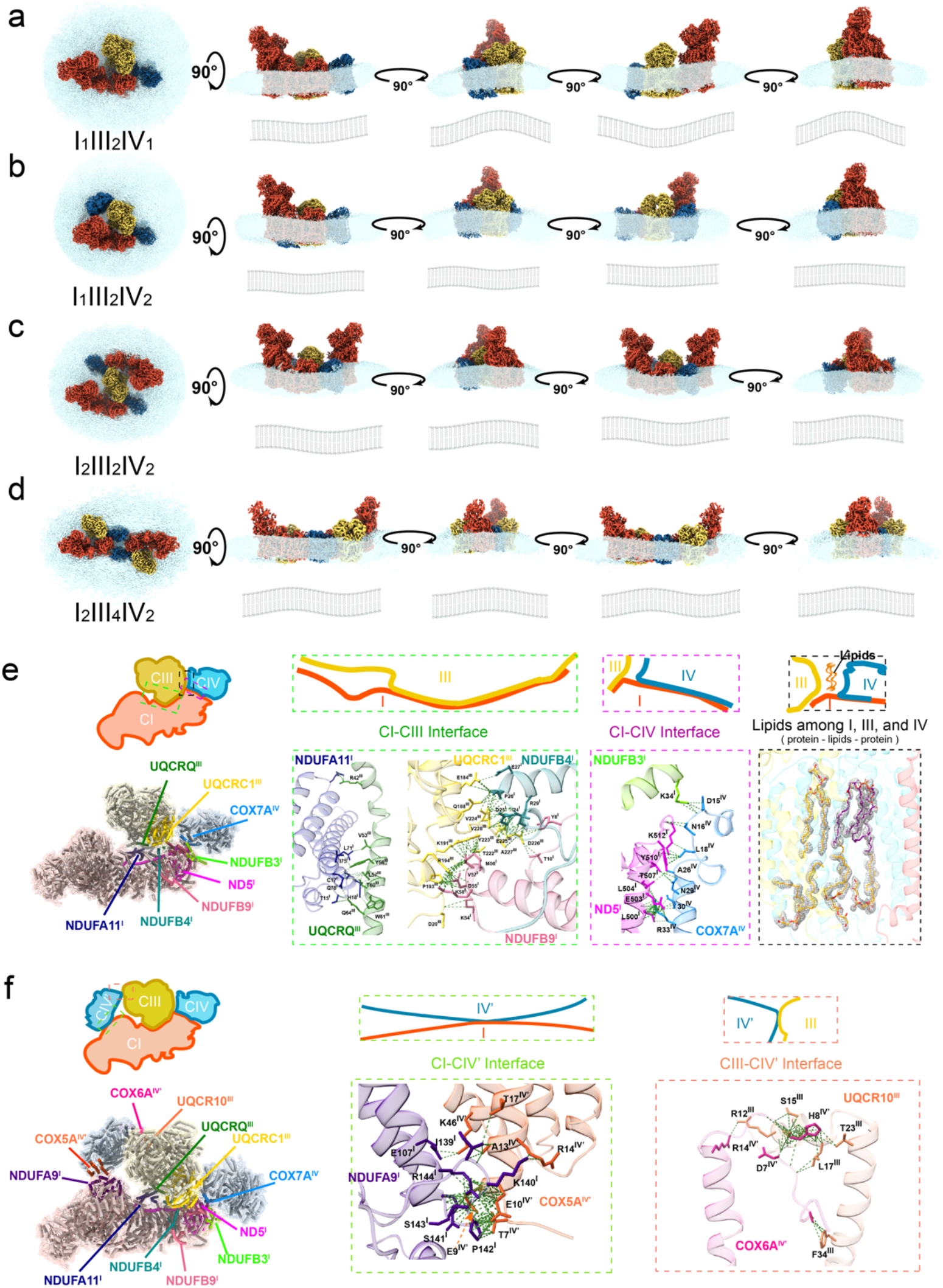
| Influence of supercomplexes on surrounding membrane curvature. **a-d**, Side views illustrating the impact of supercomplexes on membrane curvature. **e,f**, Atomic-level details of interaction interface among individual complexes. The interactions involve both protein-protein and protein-lipids-protein interactions (Supplementary Video 3). In particular, no direct protein-protein interactions are observed between CIII and CIV. Dashed lines indicate close atomic contacts with van der Waals (VDW) overlap greater than 0.4 Å, analyzed in ChimeraX.

**Extended Data Fig. 11.**
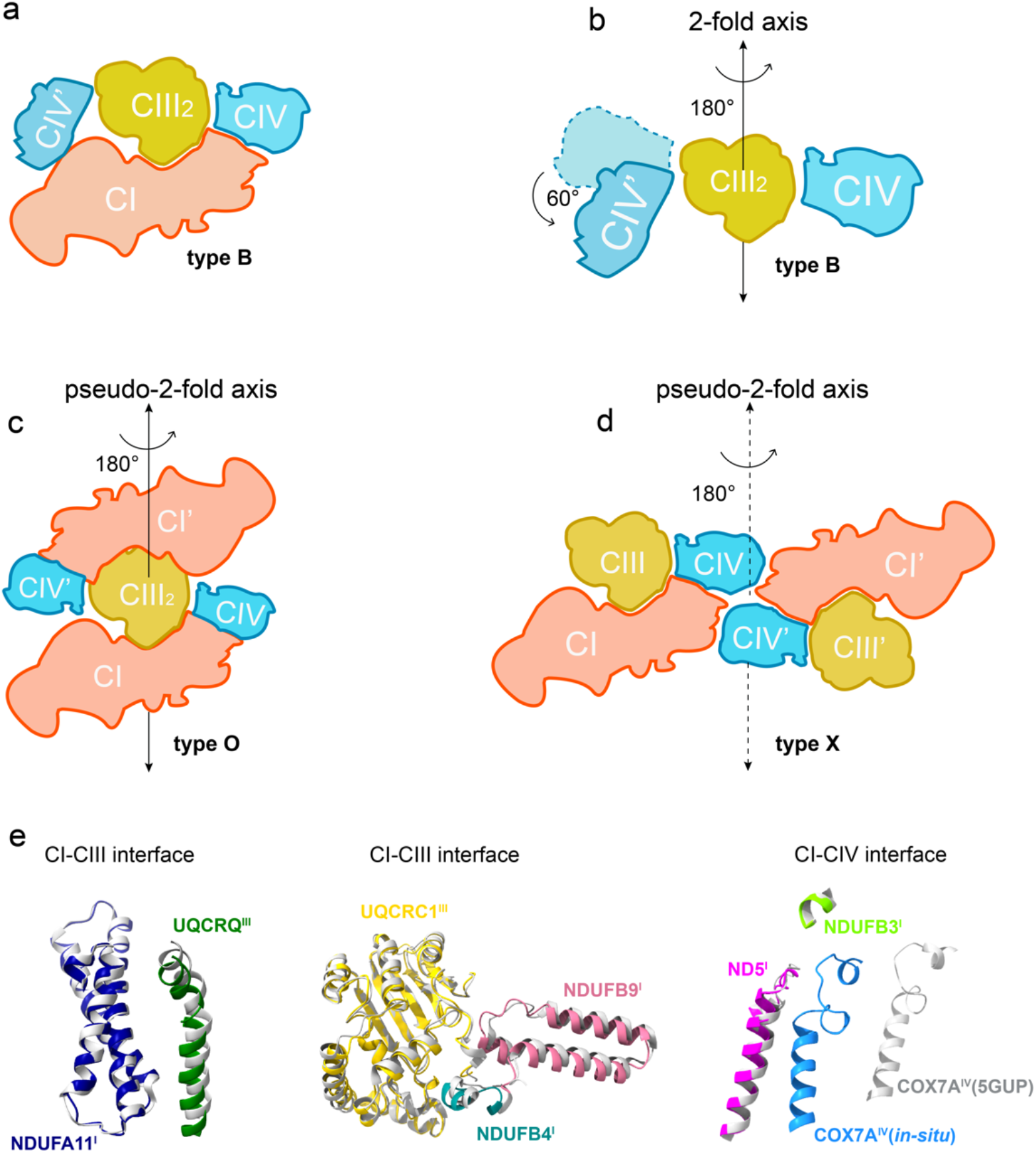
| Architecture and symmetry of *in-situ* supercomplex structures and interface comparison with the *in-vitro* supercomplex I_1_III_2_IV_1_. **a,b**, Type B supercomplex exhibits asymmetry in its two CIVs along the 2-fold axis of the CIII dimer. **c,d**, Both type O and type X supercomplexes display a pseudo-2-fold axis due to membrane curvature-induced distortion. **e**, Interface comparison between individual complexes CI-CIII and CI-CV in our *in-situ* type A supercomplex (colored) and the *in-vitro* supercomplex I_1_III_2_IV_1_ (white, PDB code: 5GUP) reveals evident differences.

**Extended Data Fig. 12.**
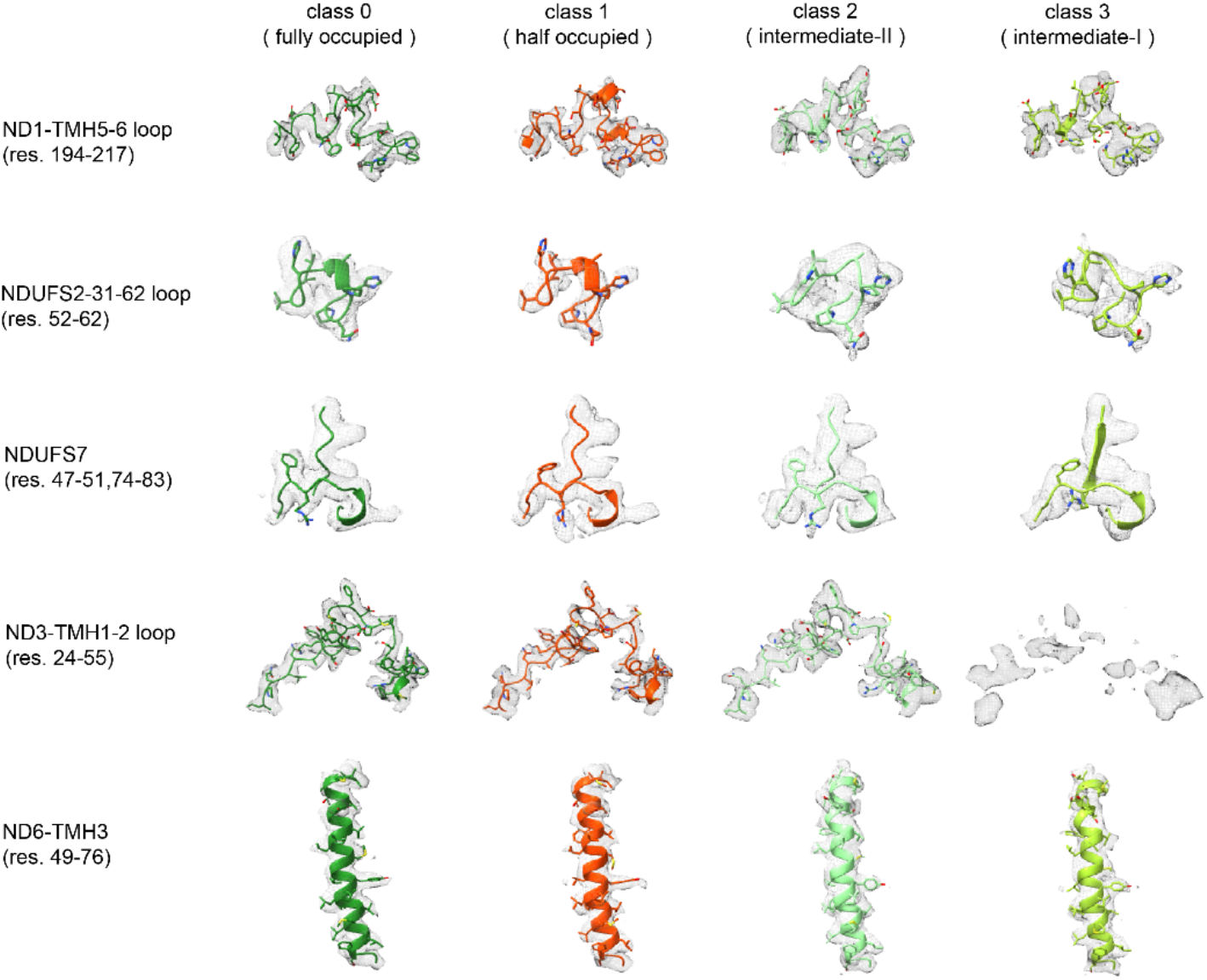
| Analysis of the hallmarks of the four distinct Q-occupied states. Class 0-2 exhibit standard active features, while class 3 show partial deactive features, suggesting these conformational changes may be required for the intermediate states during the reaction.

**Extended Data Fig. 13.**
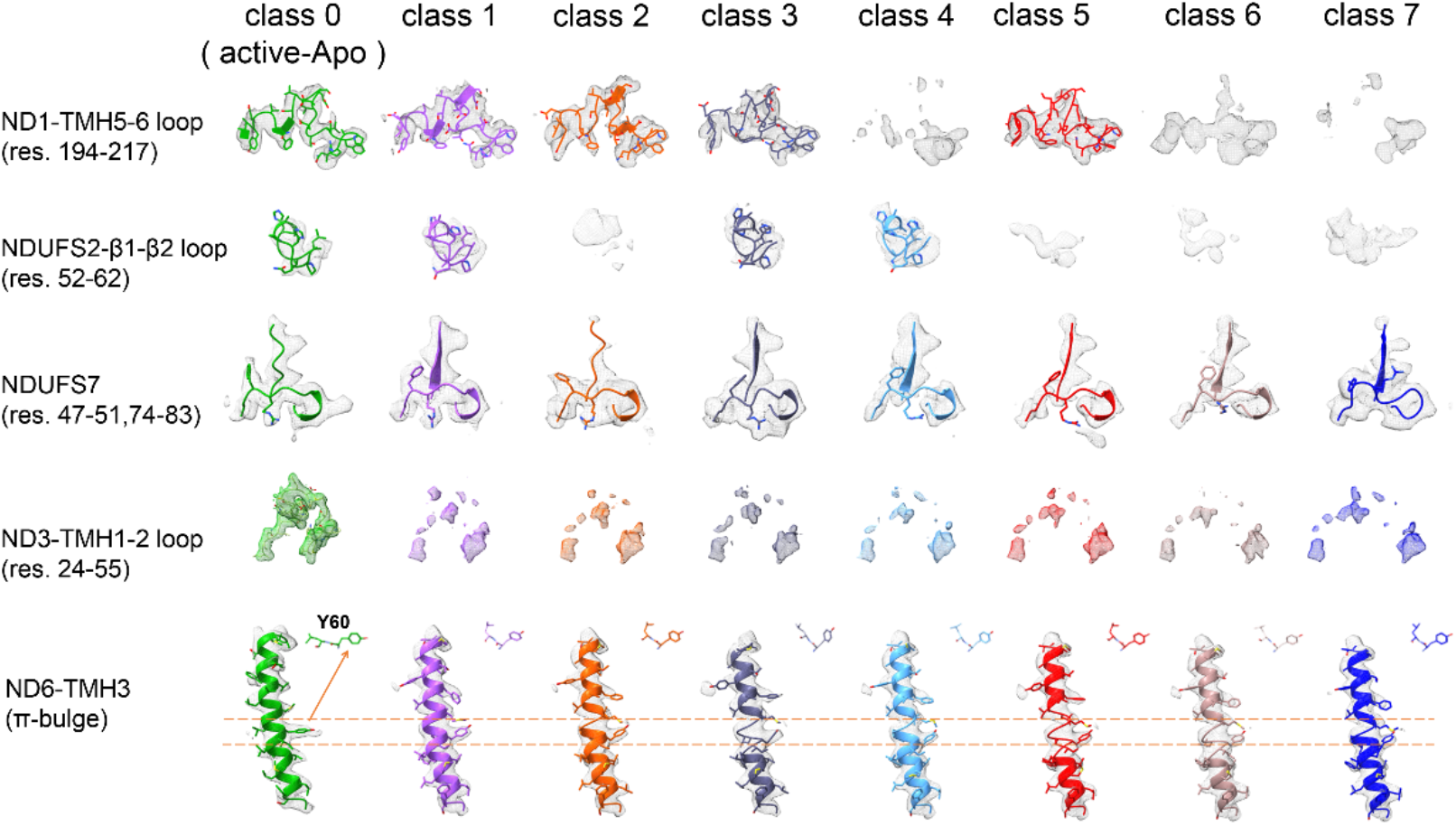
| Focused 3D classification targeting the Q-site identifies 7 major intermediate classes. Two extreme classes (class-0 and 7) fully conform to all the hallmarks of the active and deactive states, respectively. Other classes exhibit only a subset of the hallmarks associated with the deactive state, suggesting that they represent various stages in the transition between the two extreme states.

**Extended Data Fig. 14.**
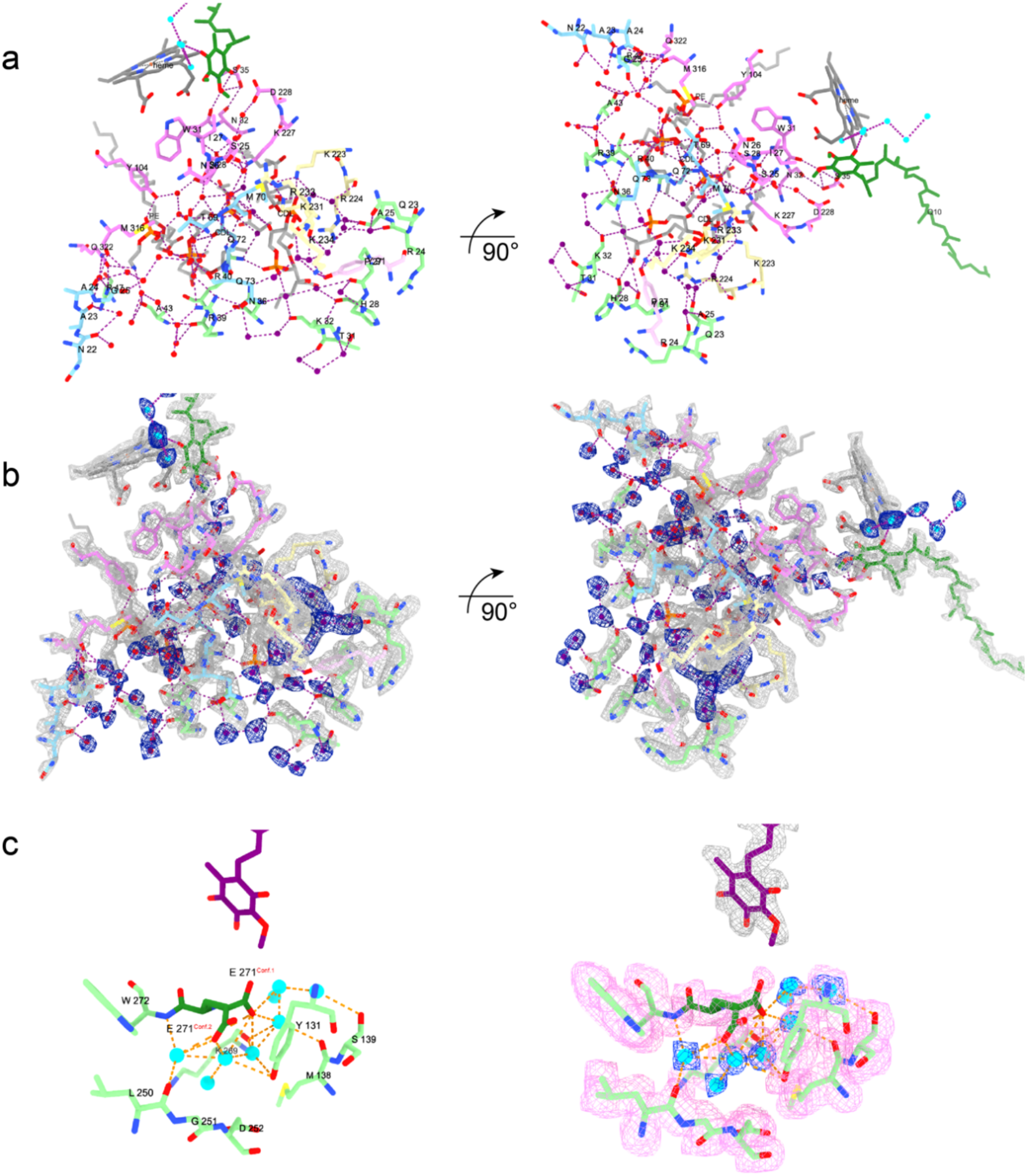
| Complex hydrogen-bonded networks near the Q_i_ sites for proton transfer in CIII. **a**, These networks consist mainly of water molecules, supported by their interacting residues, and polar headgroups of lipids, forming the proton uptake path near the Q_i_ site and the proton release path near the Q_i_ site. **b**, The high-resolution map enables us to confidently build water molecules in Q_i_ site and their surrounding residues and lipids that form these intricate hydrogen-bonded networks. **c**, The proton-transfer water chain and local density map near the Q_o_ Site.

**Supplementary Table 1.**
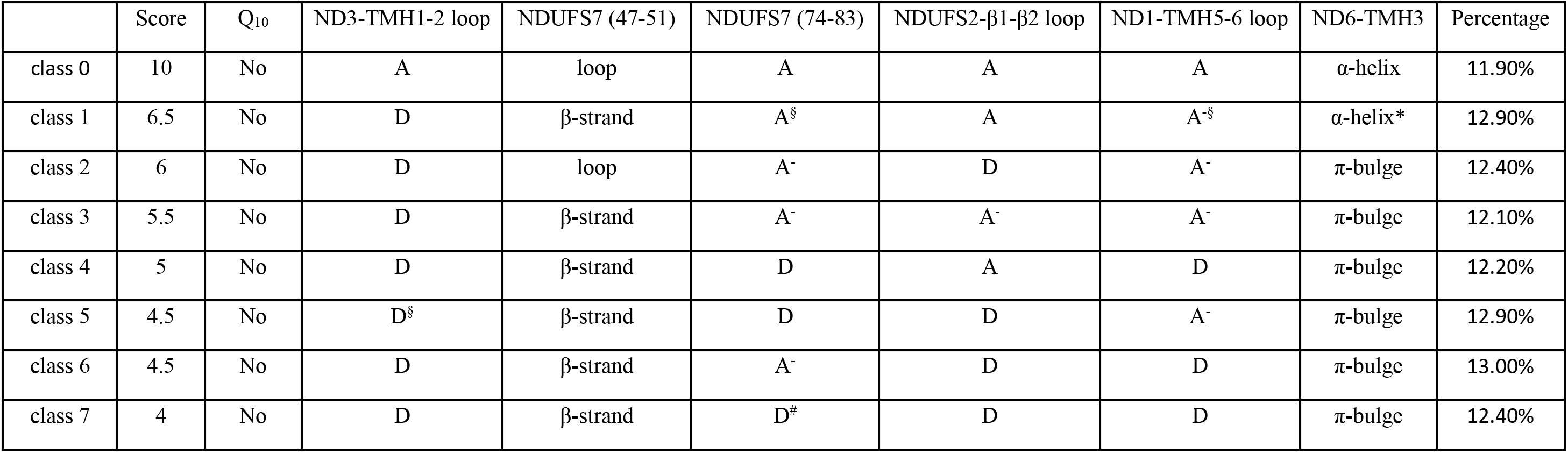
| Analysis of “hallmarks” across different classes. § D denotes the confidently identified deactive state feature, while A represents the confidently identified active state feature. A^-^ indicates that the active state feature remains recognizable but shows some changes. # Arg77 and Phe76 shift and obstruct the Q_10_ channel. Although it is still recognized as an α-helix by ChimeraX, the conformation notably differs from the active state and closely resembles a π-bulge. The various states were scored based on the hallmarks. The fully active state was assigned 10 points. If it exhibited deactive features, 1 point was deducted. If it fell between the active and deactive states, 0.5 points were deducted. Class 0 represents a fully active apo state, while class 7 is categorized as the fully deactive state.

